# ABEL-FRET: tether-free single-molecule FRET with hydrodynamic profiling

**DOI:** 10.1101/786897

**Authors:** Hugh Wilson, Quan Wang

**Affiliations:** Lewis-Sigler Institute for Integrative Genomics, Princeton University, Princeton, NJ 08544

## Abstract

Single-molecule Förster resonance energy transfer (smFRET) has become a versatile and widespread method to probe nanoscale conformation and dynamics. However, current experimental protocols often resort to molecule immobilization for long observation times and rarely approach the resolution limit of FRET-based nanoscale metrology. Here we present ABEL-FRET, an immobilization-free platform for smFRET measurements with near shot-noise limited, Angstrom-level resolution in FRET efficiency. Furthermore, ABEL-FRET naturally integrates hydrodynamic profiling, which harnesses single-molecule diffusion coefficient to enhance FRET sensing of biological processes.

## Introduction

Since the pioneering work by Ha et al^1^, single-molecule Förster resonance energy transfer (smFRET) has evolved into a powerful tool for probing biology at the individual molecule level^2^. By reporting nanometer distance information between two fluorescent probes, smFRET is able to peek into dynamic biological systems in real time, revealing deep mechanistic insights that are otherwise masked by ensemble averaging in traditional biochemical assays. Applications of this technology have rapidly permeated to a broad array of topics in physical chemistry, biophysics, molecular and structural biology (see the recent review by Lerner et al. ^3^ and the references within).

The majority of smFRET studies are carried out by attaching biomolecules to a surface ^4^. However, this is not always feasible or desirable, as the practice requires engineering recognition sites at the biomolecule-coverslip interface^5^ and in certain cases can influence^6^ or even inactivate^7^ biological function. Moreover, tethering to a surface prohibits the extraction of useful information from the diffusive motion of single molecules^8^. On the other hand, conventional tether-free smFRET measurements^9^ are limited to ~1ms observation time per molecule, often too short for biological dynamics. These limitations motivated recent efforts to develop tether-free smFRET modalities with long observation times^10,11^ (see also Ref. ^12^ and the references therein).

We present ABEL-FRET, a platform that combines high-precision smFRET spectroscopy with the ability to isolate single molecules in solution by Anti-Brownian ELectrokinetic (ABEL) trapping^13,14^. Though the idea of combining smFRET with the ABEL trap has long been proposed^15^, early experimental attempts^16^ were limited by technical difficulties, which were recently overcome^17^. With ABEL-FRET, we demonstrate continuous smFRET recording in solution over several seconds without immobilization. Compared with existing smFRET modalities, ABEL-FRET achieves unprecedented precision in FRET efficiency, resolves dynamics in a wide range of biological contexts, excels in several other performance metrics (Table S2), and adds single-molecule diffusion coefficient as a new observable to sense folding and binding stoichiometry along with molecular conformation (i.e. hydrodynamic profiling).

## Results

### ABEL-FRET implementation and example trace

ABEL-FRET combines a feedback trapping apparatus with smFRET detection optics. To trap a single molecule, the ABEL trap monitors its position in real time and applies feedback voltages to approximately cancel Brownian motion via electrokinetic forces^13^. We implemented an advanced version of the ABEL trap using rapid beam scanning, optimal real-time signal processing and a fused silica microfluidic sample holder^17^ (Figure 1a, S1 and S2, Supplementary Method). This design encodes position information in the arrival time of individual photons and achieves the ultimate feedback bandwidth limited by the fluorescence detection rate (~10-50 kHz). Similar implementations have previously demonstrated trapping of small (<5nm) biomolecules for 10s of seconds and even single fluorophores (<1nm) for a couple of seconds^18,19^. To enable smFRET on the ABEL trap platform, we implemented dual-channel photon counting and green (532nm) excitation that can be switched between continuous-wave and pulsed modes (Figure S1). In the continuous-wave mode, ratiometric photon counting in the donor and acceptor detection channels yields the smFRET efficiency. In the pulsed excitation mode, time-correlated single-photon counting was employed to monitor smFRET via a reduction of donor excited-state lifetime. To ensure trapping with maximum efficacy, donor and acceptor photons are combined on the control hardware to generate feedback actuation (Figure S2).

**Figure 1.**
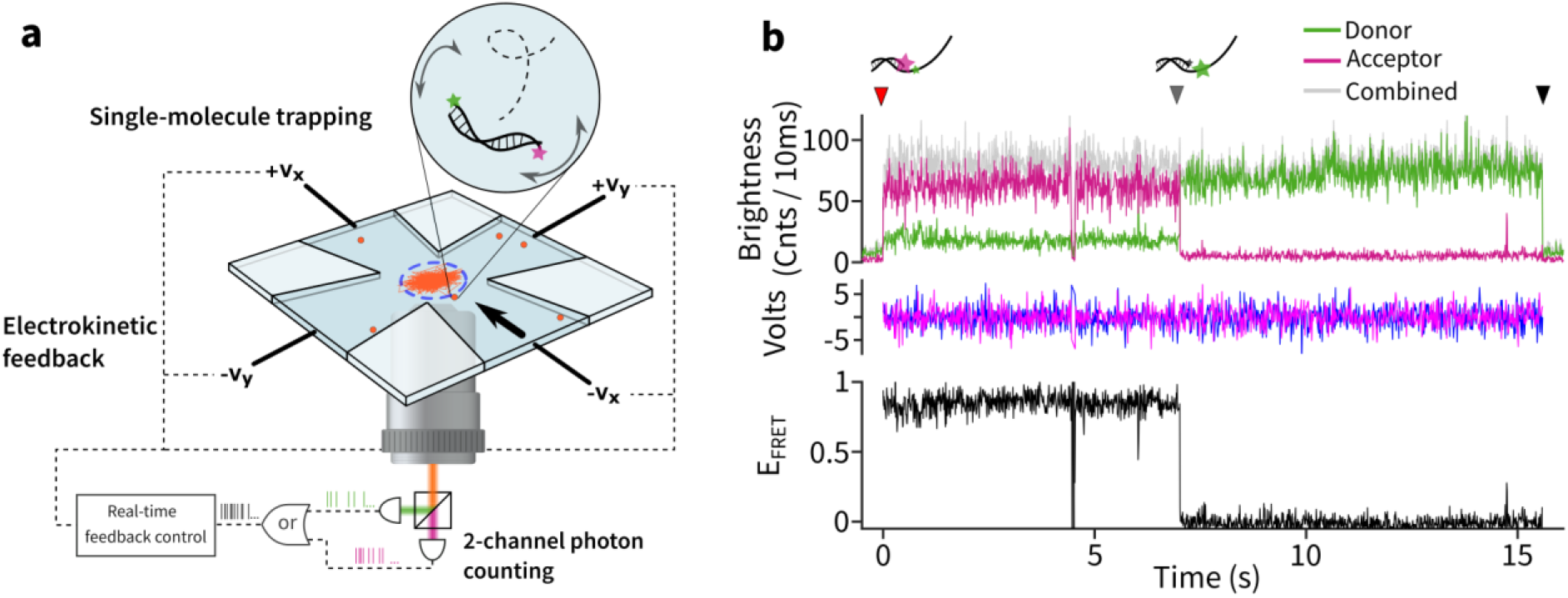
Schematic and example measurement of ABEL-FRET. (**a**) Overview of ABEL-FRET instrumentation. FRET-labeled single molecules (orange dots) are freely diffusing in a microfluidic device. Once a single molecule enters the trapping region (blue dotted circle), it is kept there by position dependent, feedback voltages that counteract Brownian motion. Fluorescence photons are separated into donor and acceptor channels and time tagged for high time resolution FRET recording. The timing of the photons also encodes position information and is used to generate accurate feedback vectors in 2D (black arrow). (**b**) An example ABEL-FRET measurement showing fluorescence brightness (green: donor, red: acceptor, grey: total intensity), feedback voltage (magenta: x, blue: y) and FRET efficiency, with 10ms binning. Red, grey and black arrows indicate molecule entry, acceptor photobleaching and trap loss events, respectively.

One example of ABEL-FRET measurements is shown in Figure 1b. Here the sample is partial duplex DNA with the donor (Cy3) and acceptor (Cy5) separated by 10 bases of single-strand DNA (ssDNA). The trace begins with no molecule in the trap (before time 0) and displays only background counts in each channel. Shortly after (indicated by a red arrow), a DNA molecule diffused into the trapping region (panel a, blue dashed circle) and was captured for ~15 seconds by the applied feedback voltages (Volt panel). While a molecule is captured, smFRET efficiency can be continuously calculated at configurable time resolutions (10ms in this example). Notably, at about 7 seconds (grey arrow) the acceptor photobleached, giving rise to an anti-correlated, step-wise change in donor and acceptor intensities. At about 16 seconds, the trapping event terminated, mostly likely due to photobleaching of the donor, freeing the trap for another molecule. Typically several hundred molecules can be measured sequentially in one hour. ABEL-FRET thus enables long-term, continuous smFRET measurement without surface tethering.

### Precise FRET efficiency measurements near the shot-noise limit

To characterize ABEL-FRET, we focus on measurement precision. The precision of a FRET efficiency measurement is fundamentally limited by the Poisson noise of photon counting (e.g. the shot-noise limit)^20,21^. Surprisingly, only very few smFRET measurements have approached this shot noise limit^22–24^. To characterize the measurement precision of ABEL-FRET, we chose short (11-13bp) double-stranded DNA molecules due to their high rigidity and distance tunability. The DNA molecules were labeled at the 3’ ends with Cy3 and Cy5 as the FRET pair and were reliably captured for multiple seconds, yielding >10,000 photons (donor and acceptor combined) per molecule (Figure 2a). After recording smFRET traces for 226 molecules (here 11bp dsDNA), we divided each trapping event into batches of *N* consecutive photons, calculated the FRET efficiency of each batch and constructed a histogram by pooling data from all molecules. We fit the resultant histogram using a Gaussian distribution and extracted the standard deviation (*σ*) to quantify measurement precision. As seen in Figure 2a, increasing the number of photons per batch from 750 to 6,000 resulted in a ~2.5 fold improvement in precision (*σ* from 0.024 to 0.0095). Next, we characterized the precision (*σ*, Figure S3) over two decades of photon numbers (*N* from 100 to 20,000) and plot the scaling behavior in Figure 2b. In parallel, we estimated the shot-noise limit both by numerical simulation (red dashed line, Figure 2b, Supplementary Note 1, Supplementary Table 1) and by deriving the Cramér-Rao bound (Supplementary Note 2, Figure S4) from a maximum likelihood framework^20^. Evidently, ABEL-FRET tracks the shot-noise limit with ~ 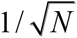 scaling and achieves an absolute value no more than 30% above the shot-noise limit.

**Figure 2.**
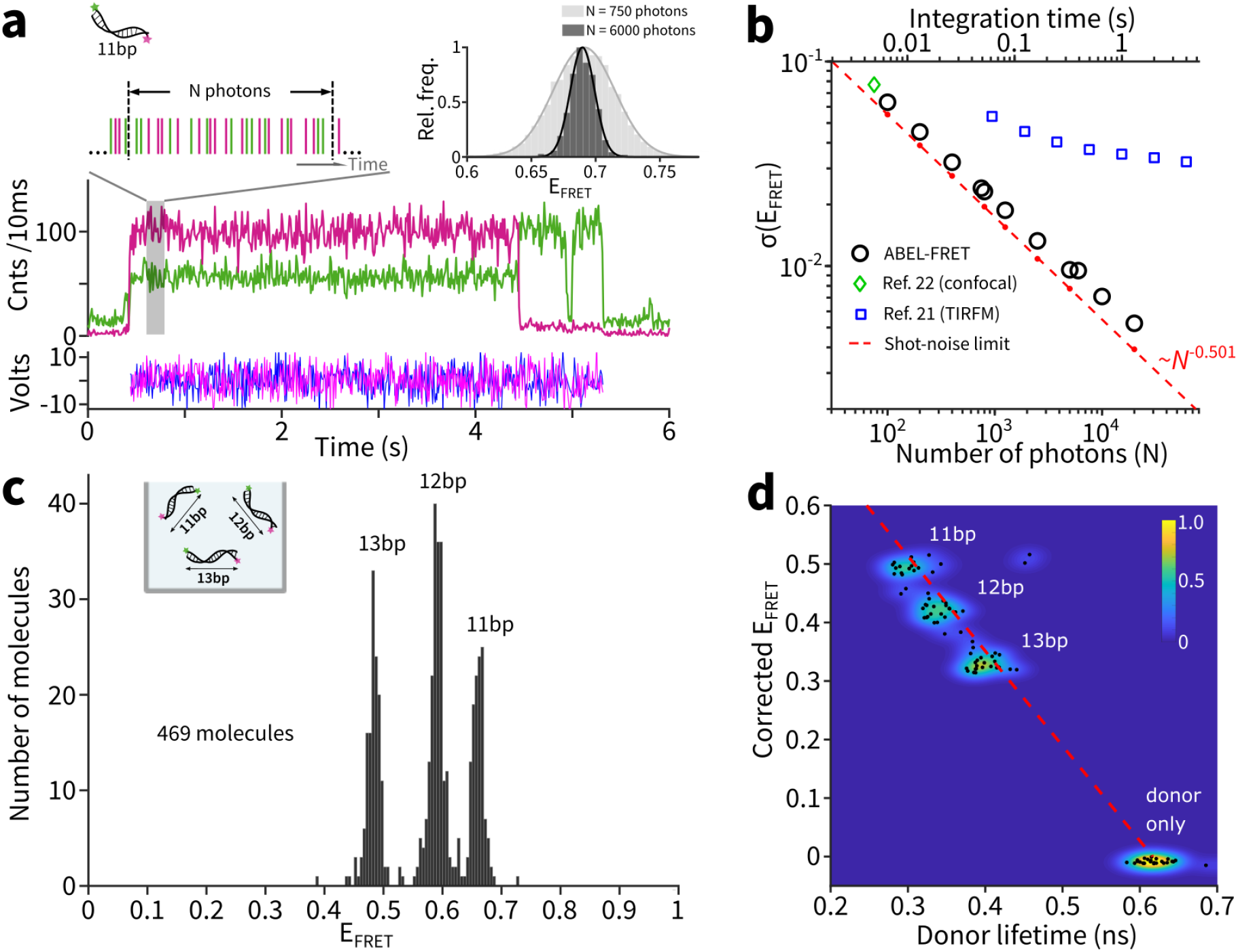
ABEL-FRET measures FRET efficiency with near shot-noise limited precision. (**a**) Scheme to characterize measurement precision using 11bp dsDNA, showing a sample trace, the definition of *N* (number of photons per batch) and example efficiency histograms. (**b**) FRET precision (σ) as a function of photon number *N*. The dotted red line is a power law fit of the simulated precision (red dots). Integration time (top axis) is calculated from *N* using a measured detection rate of 15.3kHz. (**c**) FRET efficiency (molecule average) histogram of a mixture sample composed of 11, 12 and 13 bp dsDNA molecules. (**d**) 2D scatter plot of FRET efficiency versus donor lifetime for the same mixture sample as in panel **c**. The underlying density is visualized using a 2D kernel density estimation algorithm. The dotted red line shows the theoretical relationship E=1-τ/τ_0_, where the value of τ_0_ is extracted from the mean lifetime of the donor only species.

Additionally we compare ABEL-FRET to popular smFRET modalities: burst-based confocal measurements (data extracted from Ref.^22^) and Total Internal Reflection Fluorescence Microscopy (TIRFM, data extracted from ^21^). As expected, the precision of burst-based confocal measurements is poor due to the low number of photons (~50) detected per molecule and agrees with the low-photon limit of ABEL-FRET. On the other hand, TIRFM is able to harvest more photons per molecule but the achievable precision deviates from the shot-noise limit and is at least two times worse than ABEL-FRET (see discussion).

### Resolving single base pair differences in dsDNA mixtures

To explore the practical limit of resolution, we challenged ABEL-FRET to resolve a mixture of 11,12 and 13bp dsDNA duplexes, all end labeled with a Cy3-Cy5 pair (Table S3). In this case, the molecules differ by only a single base pair, which corresponds to ~0.33nm rise along the helical axis. Such a small difference was at the resolution limit in previous smFRET attempts^21^. We trapped molecules one-by-one and histogrammed the mean FRET efficiency of 469 molecules (Figure 2c). ABEL-FRET clearly resolves the three populations (Figure S6). It is well known that terminally attached cyanine dyes tend to stack at the end of the helix^25^. We thus interpret the observed differences in FRET efficiency to have contributions from both distance and orientation changes along the helix.

We next switched to pulsed excitation and simultaneously measured donor lifetime and FRET efficiency of individual molecules of the mixture (Figure S7). This allows us to build a two-dimensional scatter plot to separate and identify populations in efficiency-lifetime space. Figure 2d confirms that 1bp differences are resolvable and the observed populations follow the expected relationship between the two variables (E=1-τ⁄τ_0_, where τ_0_ is the donor lifetime without the acceptor). This measurement demonstrates ABEL-FRET’s compatibility with excited-state lifetime measurements^26^, which provide complementary information on nanoscale dynamics^27^.

### Tether-free smFRET dynamics of biomolecules from millisecond to second

We next demonstrate continuous monitoring of smFRET dynamics of biomolecules without tethering and compare to previous surface immobilized measurements. First, we measured the spontaneous interconversion between the two stacked isomers of the Holliday junction (HJ) ^28^ (Table S3). With ABEL-FRET transitions between two FRET levels were clearly visualized with 50mM (Figure S8) and 5mM Mg^2+^ (Figure 3a and S8). Using cross-correlation analysis^29^ (Figure 3b), ABEL-FRET easily resolves kinetic rates from ~10 s^−1^ (50mM Mg^2+^) to ~500 s^−1^ (1mM Mg^2+^, Figure S8). Comparing to measurements on immobilized molecules^28^, we found excellent agreements in kinetic rates (Figure 3c), the efficiency values (~0.6 and ~0.2) as well as the relative abundances of the two conformers (Figure S9). Furthermore, the kinetic rates are independent of the feedback strength (Figure S11). These results eliminate the concern that the feedback electric fields might perturb HJ behavior (Supplementary Note 3).

**Figure 3.**
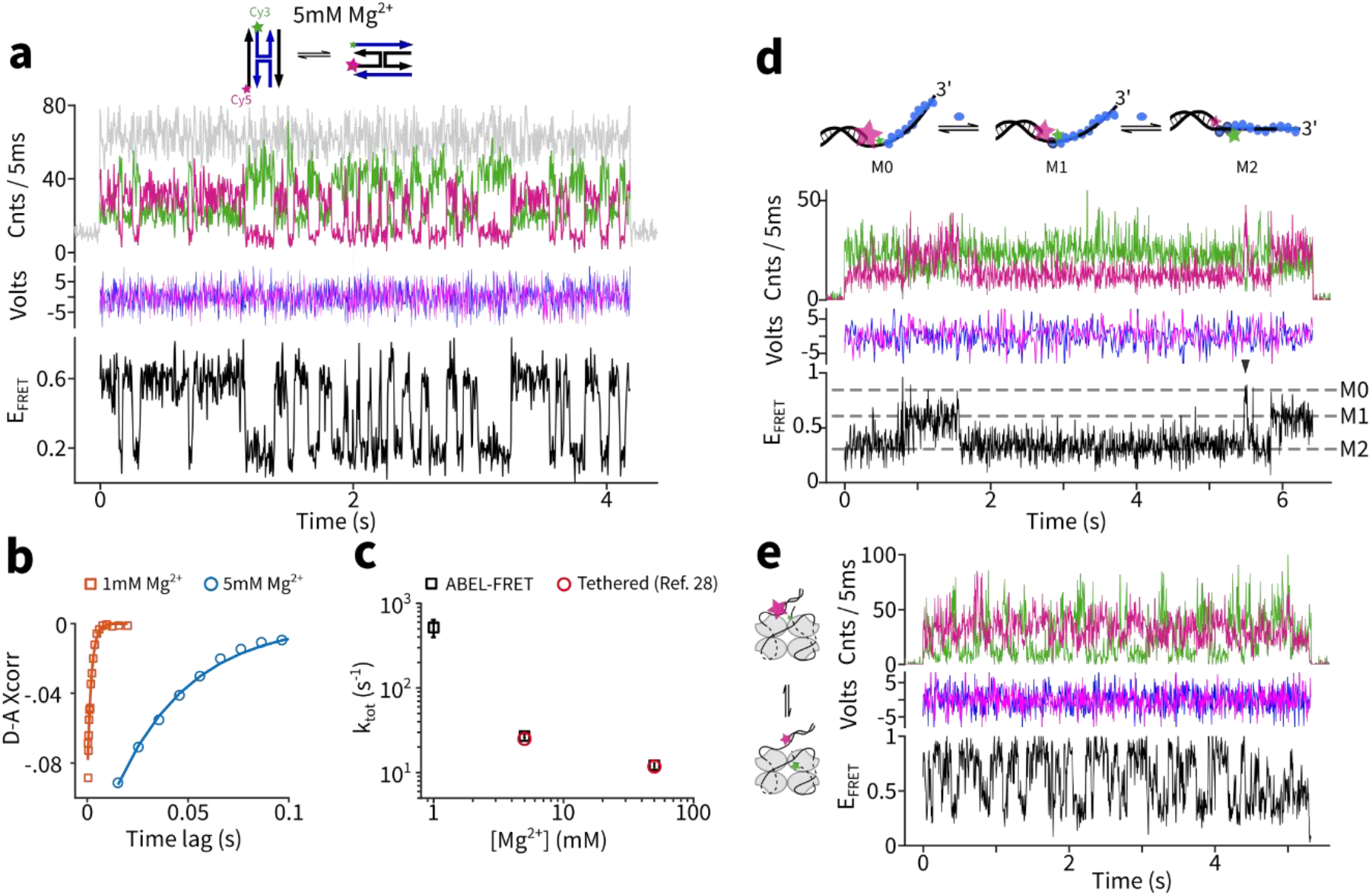
Tether-free recording of biomolecular dynamics from millisecond to second timescales. (**a**) Example trace of a single Holliday Junction (HJ) with 5mM Mg^2+^ including fluorescence intensity (green: donor, red: acceptor, grey: total intensity), feedback voltages and FRET efficiency. (**b**) Donor-acceptor intensity cross-correlation curves of HJ for 1mM Mg^2+^ (orange, averaged over 44 molecules) and 5mM Mg^2+^ (blue, averaged over 59 molecules). Solid lines are single exponential fits. (**c**) Extracted rates of HJ dynamics and comparison to literature values. Error bars represent 95% C.I. (**d**) Top cartoon: RecA protein (blue oval) binding and unbinding in the ssDNA region between the donor and acceptor leads to FRET changes. Bottom: an example ABEL-FRET trace with 1µM RecA, 2mM ATP. Black arrow indicates a transient event. (**e**) Left cartoon: SSB protein sliding on ssDNA causes rapid FRET fluctuations. Right: an example trace with 0.5nM SSB, 200mM NaCl.

We then measured two model systems from DNA metabolism in bacteria^30^: RecA recombinase and single-stranded DNA-binding protein (SSB). Both proteins bind to ssDNA but organize their substrate in distinct, dynamic ways. To probe RecA-ssDNA dynamics free in solution, we used a Cy3-Cy5 labeled partial duplex as the trapping target^31^ (Table S3), and observed transitions between three FRET states (M0~0.87, M1~0.58, M2~0.35, Figure 3d and S12) in the presence of RecA (1µM) and ATP (2mM). Those observed FRET states and dynamics are similar to TIRFM measurements on immobilized DNA substrates^31^ and were previously interpreted to represent RecA protein binding and unbinding. Notably, our FRET efficiency histogram (Figure S12) is much sharper compared to that obtained on immobilized molecules^31^ and hints the existence of more states (i.e. E~0.45). We also observed many transient (<100ms) states (arrow in Figure 3d) which were likely missed in previous TIRFM measurements^31^. These observations highlight the enhanced precision and time resolution of ABEL-FRET and suggest extra complexity in RecA nucleofilament dynamics.

We next measured single-molecule SSB-ssDNA interactions without surface immobilization. SSB protein is known to interact with ssDNA with a footprint of either 35 or 65 nucleotides ((SSB)_35_ and (SSB)_65_ modes) and is highly mobile when bound to ssDNA substrates^32^. Using a Cy3-Cy5 labeled substrate (Table S3) and under the condition that favors (SSB)_65_ formation (200mM NaCl, 500pM SSB), we successfully recapitulated the rapid FRET fluctuations produced by (SSB)_65_ sliding on ssDNA^33^ (Figure 3e and S13), with a rate (16.9±0.9 s^−1^, Figure S13) that is consistent with the previous TIRFM measurement (~15s^−1^, interpolated from Ref. ^33^). With these examples, we demonstrate ABEL-FRET’s compatibility with a wide range of biological processes.

### ABEL-FRET enhances smFRET with hydrodynamic profiling

Finally, we demonstrate hydrodynamic profiling: using single-molecule diffusion coefficient inference^34^, together with smFRET measurement, to probe a two-dimensional space of molecular size and conformation. As a first example, we probed the Mg^2+^ induced folding of the HJ. Without Mg^2+^, most HJs exhibit a constant FRET efficiency of ~0.3 (Figure 4a), a behavior that deviates from the two-state toggling behavior observed in the presence of 50mM Mg^2+^ (Figure 4b). With hydrodynamic profiling, we get more structural information that illuminates this difference. More specifically, we extracted the diffusion coefficient (D) of every observed FRET level (Figure 4a and 4b, solid lines) and plot all single-molecule data on the D-FRET parameter space (Figure 4c). Here, FRET efficiency probes the one-dimensional distance between the two labeled positions and D reflects the global compactness of the HJ molecules. Strikingly, the ~0.3 FRET state, observed without Mg^2+^, is associated with a lower diffusion coefficient (70.2±0.7 µm^2^/s) while the two Mg^2+^-induced isomers have identical and higher diffusion coefficient (high FRET state: 81.3±0.9 µm^2^/s and low FRET state: 80.9±1.4 µm^2^/s, Figure S15). These data provide direct evidence that the HJ exists in an extended conformation without divalent ions and folds into two interconverting isomers with Mg^2+^, as previously inferred from comparative gel electrophoresis^35^.

**Figure 4.**
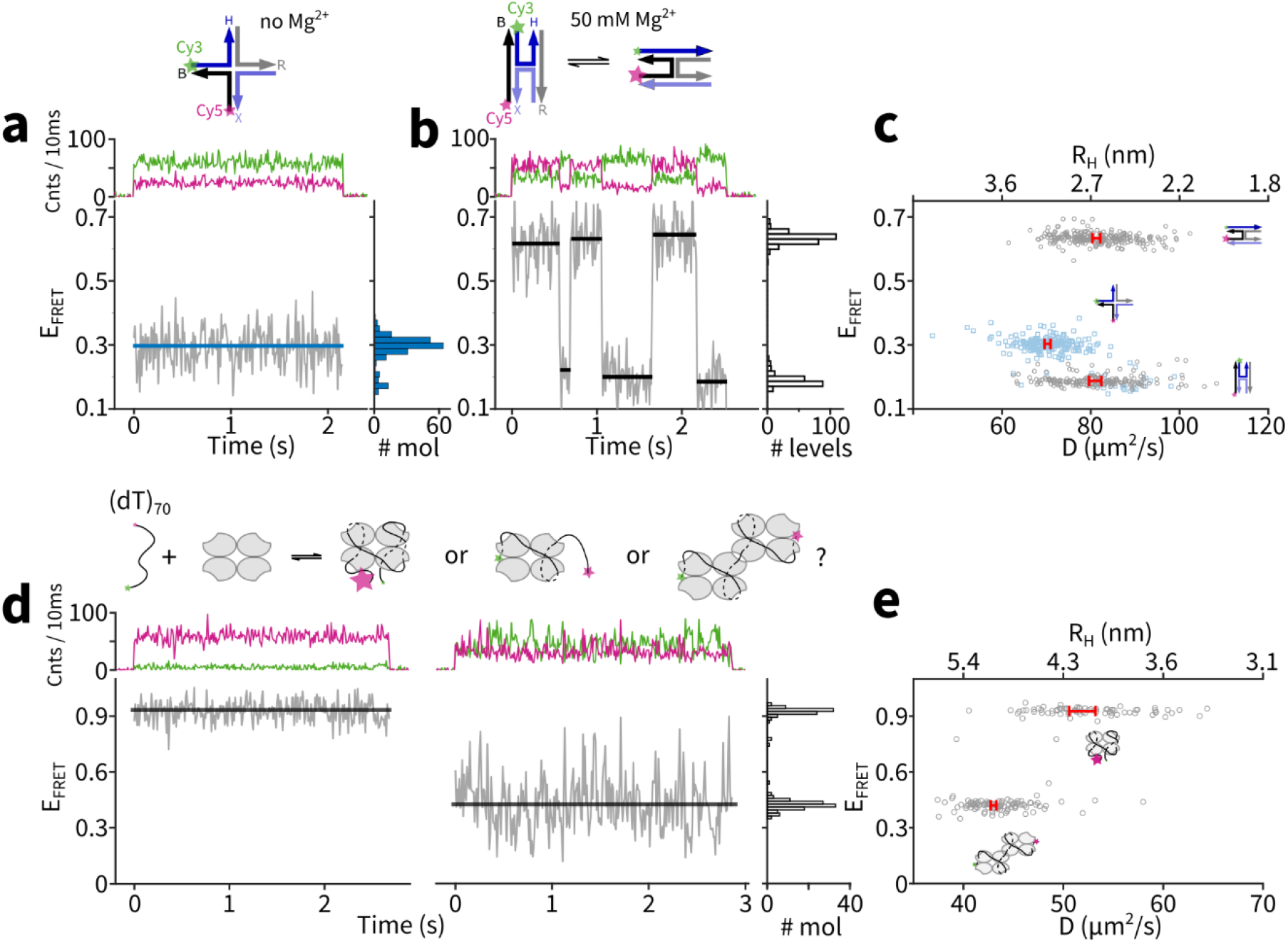
ABEL-FRET enhances smFRET with hydrodynamic profiling. (**a-c**) Probing Mg^2+^ induced folding of the HJ (**a**) Left: example ABEL-FRET trace of a single HJ with no Mg^2+^. Solid blue line represents mean FRET efficiency of the molecule. Right: FRET histogram from 196 molecules. (**b**) Left: example trace of a single HJ with 50mM Mg^2+^. Solid black lines represent identified FRET levels. Right: FRET histogram of discrete levels (439 levels from 226 molecules) (**c**) Scatter plot of the diffusion coefficient (D) and FRET efficiency of identified FRET levels in **a** and **b**. Light blue symbol: no Mg^2+^, grey symbol: 50mM Mg^2+^. Red intervals indicate the uncertainties (95% C.I.) in average D values of the respective populations. The hydrodynamic radii (R_H_) are calculated from the diffusion coefficients using the Stokes-Einstein equation. (**d-e**) Probing binding stoichiometry of SSB to ssDNA. (**d**) Top cartoon: three possible configurations of SSB-ssDNA complex. Bottom: two representative ABEL-FRET traces and FRET histogram with 10nM SSB and 40mM NaCl (186 molecules). Solid black lines are molecular averages of FRET efficiency. (**e**) Scatter plot of the diffusion coefficient (D) and FRET efficiency of all single molecules probed in panel **d**. Red intervals indicate the uncertainties (95% C.I.) in average D (or R_H_) values of the respective populations.

As a second example, we use hydrodynamic profiling to determine the binding stoichiometry of SSB-ssDNA complexes^36^. On an end-labeled 70nt ssDNA substrate, three binding scenarios are possible (Figure 4d, top cartoon): one SSB tetramer fully wrapped with a footprint of 65 bases (high FRET), one SSB tetramer partially wrapped with a footprint of 35 bases (low FRET), or two SSB tetramers fully wrapped with each occupying 35 bases (low FRET). Note that in this case FRET reports the end-to-end distance of the ssDNA, not directly on SSB binding stoichiometry. On the other hand, the diffusion coefficient directly reflects the number of SSB tetramers bound. Here, ABEL-FRET measurements identified a bimodal distribution of FRET populations (Figure 4d and S14), and through hydrodynamic profiling (Figure 4e and S15), revealed that the low FRET state diffuses slower (E~0.45, D_mean_=43.0±0.3 µm^2^/s) than the high FRET state (E~0.93, D_mean_=51.9±1.3 µm^2^/s). This difference can be fully accounted for by the hydrodynamic volume of a second SSB tetramer (Supplementary Note 4). We thus provided direct evidence that the SSB-ssDNA (70nt) complex contains either one SSB tetramer in the 65nt binding mode, or two SSB tetramers in the 35nt binding mode.

## Discussion

Here we have introduced ABEL-FRET, which employs feedback trapping to allow continuous observation of smFRET on freely moving and rotating molecules in solution, without the need to immobilize or encapsulate the target of interest. ABEL-FRET maintains a practical throughput (~600 mol/h), naturally integrates excited-state lifetime measurements^26,27^, features ~ms time resolution, is compatible with common biochemical buffers and should work on any FRET-labeled biological molecules. A detailed comparison to other smFRET modalities can be found in Table S2. ABEL-FRET could be further enhanced by incorporating alternating excitation^37^ or non-equilibrium measurements^38,39^.

ABEL-FRET achieves a precision in FRET efficiency that approaches the fundamental limit imposed by the Poisson statistics of photon counting. In Figure 2c, the standard deviation of a single population is ~0.009, which corresponds to an ultimate resolution of ~0.006R_0_ (~30pm near R_0_=5nm) in distance space. This level of precision in smFRET spectroscopy has not been achieved before and will realize the full potential of smFRET as a molecular ruler for biological application. Although nothing fundamental prevents current smFRET technology from reaching the Angstrom-level precision demonstrated here, the act of immobilization might introduce extra heterogeneity that broadens FRET distributions, as long been speculated in the field^21,40^. ABEL-FRET, given its complete elimination of potential tether-induced effects, will allow researchers to focus on the intrinsic heterogeneity between molecules.

ABEL-FRET is the only platform that measures hydrodynamic size and smFRET simultaneously. By combining these two orthogonal degrees of freedom, ABEL-FRET opens up new possibilities in single-molecule sensing, particularly for biological processes of higher complexity^41^. Due to its tether-free sample preparation, shot noise limited precision at the Angstrom level and hydrodynamic profiling capability, ABEL-FRET represents a major technology upgrade that will benefit all smFRET measurements.

## Online Methods

### ABEL-FRET setup

The ABEL trap was implemented based on a published design^42^ (Figure S1). Briefly, light from a 532nm laser (Coherent Obis) was scanned by a pair of acousto-optic deflectors (AA Opto-electronic) to produce a 32-point ‘knight’s tour’ pattern at the sample plane (~3 µm × 3µm). The scan speed was set to 600ns per point, which achieves an effective imaging frame rate of ~52kHz. The molecule’s real-time position was estimated on a Field Programmable Gate Array (FPGA, National Instruments PCIe-7852R) with photon-by-photon mapping and a Kalman filter^43^ (Figure S2). Feedback voltage vectors were calculated using 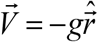 where 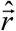 is the filtered position estimate and *g* is a gain parameter (volt per µm of displacement). The voltages were applied to the microfluidic chip using a pair of 20× amplifiers (FLC F10AD) and platinum electrodes. The electric field strength in the trap was estimated assuming the voltage drops uniformly over a 100 µm region. The smFRET signal collected by a silicone oil immersion objective (Olympus UPLSAPO100XS, NA 1.35) was first focused through a confocal pinhole (400 µm, Thorlabs) and then spectrally separated into donor and acceptor paths using a dichroic beamsplitter (Semrock FF652-Di01-25×36). The pass bands of the donor and acceptor channels were further narrowed by emission filters before photons were time-tagged by avalanche photodiodes (Laser Component Count T-100). The digital pulses generated by the two detectors were combined (logical ‘or’ operation) on the FPGA (12.5ns resolution) before the position estimation step. This ensures that both donor and acceptor photons were used for feedback.

To measure the time-resolved decay of the donor, excitation was provided by the spectrally-filtered (532±5nm) supercontinuum fiber^44^ (OFS Fitel) pumped by a mode-locked Ti:Sapphire oscillator (Mira 900, Coherent, tuned to 790nm). Time-correlated single-photon counting was performed using a PicoHarp 300 module (PicoQuant). Excited-state lifetime was extracted using a maximum likelihood estimator as previously described^19^. The average excitation intensity at the sample was between 150-600Wcm^−2^.

### Microfluidic device fabrication, cleaning and passivation

The microfluidic devices used for ABEL-FRET experiments were fabricated in Princeton University’s Micro/Nano Fabrication Laboratory (MNFL) using a protocol modified from those published^45,46^ (Supplementary Methods). After the devices were made, they were cleaned and passivated with poly(ethylene glycol) (PEG) using the following protocol. First, the device was cleaned by an overnight Piranha bath (3:1 mixture of sulfuric acid and hydrogen peroxide) followed by extensive rinsing with ultrapure water (18.2 MΩ). Next, the chip was incubated in 1M potassium hydroxide for 15 minutes. Then, the interior of the chip was filled with mPEG-silane solution (Gelest SIM6492.73 or Laysan Bio MPEG-SIL-5000-1g, >20mg/ml in 95% ethanol-5% water mixture with pH ~5)^47^ and incubated for more than 24 hours at room temperature. Finally, the chip was dried with pure nitrogen, flushed and stored in ultrapure water before use.

### Sample preparation

Sequences of DNA samples are listed in Table S3. All strands were purchased from IDT, purified by HPLC. For those that were labeled in house, the amine-modified strand was incubated with ~10 fold excess of NHS-functionalized dye (sulfonated Cyanine3 or sulfonated Cyanine5, Lumiprobe) in 0.2M sodium bicarbonate buffer pH8.3 for >4 hours. The dye-labeled oligo was then purified using ethanol precipitation (3 times) or size-exclusion columns (BioRad, P6, 2 times). The labeling efficiencies, determined by absorption measurements, were 70%-90%. Duplex DNA was annealed by mixing the two complementary strands at ~10µM concentration in 20mM HEPES pH8 100mM NaCl, heating to 95 ºC for two minutes and slowly cooling (~4 hours) to room temperature. Holliday junctions were annealed similarly, in 25mM HEPES, 50mM NaCl pH7.5. The DNA samples were stored at −20 ºC before use.

All ABEL-FRET experiments were performed at ~5pM of the labeled species in a trapping buffer containing 20mM HEPES, 3mM Trolox, and an enzymatic oxygen scavenger system (50nM protocatechuate-3,4-dioxygenase and 2.5mM protocatechuic acid). Additional components were added depending on the experiments. dsDNA: 100mM NaCl; Holliday junction: 1, 5, or 50mM MgCl_2_ or 0.5mM EDTA (no Mg^2+^); RecA: 100mM NaCl, 10mM MgCl_2_, 1µM RecA (New England BioLabs), 2mM ATP or 1mM ATPγS; (SSB)_65_ sliding: 200mM NaCl, 0.5nM SSB (Promega); SSB binding stoichiometry: 10, 40 or 100mM NaCl, 10nM SSB. Sources of all reagents are listed in Table S4.

### Data analysis

All data analysis was performed with customized software written in Matlab. First, data regions representing single molecules and background were extracted with the guide of a change-point finding algorithm^19,48^. The mean background rates for the donor, acceptor and combined channels were then determined by fitting a Poisson distribution to the intensity histogram from the background regions. The apparent FRET efficiency of trapped molecules over time was determined with background and donor leakage correction, for every time bin or batch of N photons, using

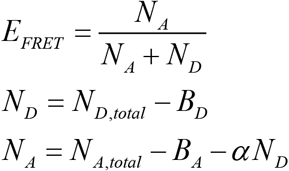

where *N*_*D*_, total and *B*_*D*_ are the total and background counts of the donor, *N*_*A*_, total and *B*_*A*_ are the total and background counts of the acceptor, α is the leakage fraction (0.074, determined by analyzing donor only dsDNA molecules). The γ correction, which takes into account the differences in quantum yield and detection efficiency between the donor and the acceptor^49^, was not included (except in Figure 2d, where it was determined from the anti-correlated steps in donor and acceptor intensity upon acceptor photobleaching using the equation γ=ΔI_A_/ΔI_D_). Molecules showing *E*_*FRET*_ < 0.05 were considered as donor only and not selected for further analysis except in Figure 2d, where the donor-only species was analyzed to extract *τ*_0_, the donor lifetime in the absence of FRET. FRET efficiency histograms were constructed on either a molecule-by-molecule basis or using a fixed number of photons as detailed in the main text. Precision was quantified by the standard deviation of a Gaussian fit to the FRET efficiency histogram. FRET state transitions in the HJ and RecA experiments are identified by a change-point finding algorithm with Gaussian statistics^50^.

The cross correlations between the donor and acceptor channels were calculated using a flexible binning algorithm^51^ based on the photon arrival times, and fit using an exponential function^29^

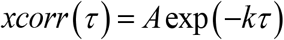

The cross correlations were calculated on single molecule data and then averaged over the molecules measured. For molecules that were trapped for more than one second, single-molecule rates were also determined.

## Acknowledgement

We thank Haw Yang and his lab members for valuable advice and feedback, Evangelos G. Gatzogiannis for the loan of a Picoharp 300 unit and the staff at Princeton’s PRISM clean room (Eric Mills, Roman Akhmechet, Zuzanna Lewicka and David Barth) for assistance with ABEL trap fabrication. This work is funded by the Lewis-Sigler Fellowship (to Q.W.) of Princeton University.

## Author Contributions

H.W. and Q.W. designed the research, performed experiments and analyzed the data. H.W. wrote an initial draft of the manuscript. Both authors discussed and interpreted results, and edited the manuscript.

