## Supplemental Information for "ABEL-FRET: tether-free single-molecule FRET with hydrodynamic profiling"

**Supplementary Methods**

**ABEL trap device fabrication procedures**

The microfluidic devices used for ABEL trap experiments were fabricated in Princeton University's Micro/Nano Fabrication Laborotory (MNFL) using a protocol modified from those published[1,2]. We chose a 4-inch, JGS1-UV grade fused silica wafer (University wafers) as the starting material due to its extremely low level of auto-fluorescence. The wafers were RCA cleaned and a uniform layer of polysilicon (~530nm, PolySi) was deposited using low pressure chemical vapor deposition (595 ºC, 3 hours). The deep channel features were first defined by direct laser writing (Heidelberg / DWL66) on spin-coated positive photoresist (AZ1518, Microchemicals GmbH). The pattern was then transferred to the PolySi layer using a $SF_6/C_4F_8$ plasma etch (Samco RIE800iPB, 100sccm $SF_6$, 100 sccm $C_4F_8$, 10W bias, 1000W ICP). After removal of the photoresist by $O_2$ plasma (TePla, 500mTorr, 500W, 25min), the deep channels with a depth of 10µm were formed by wet etching (49% HF, 8min) into fused silica using PolySi as the mask. The shallow trapping regions were subsequently formed by a second round of lithography. First, a thick (~5 µm) layer of positive photoresist (AZ 4330, Microchemicals GmbH) was spun on the remaining PolySi layer and the pattern of the trapping region was defined by contact lithography (Karl Suess MA6, 405nm, 80 seconds, 2.0mW/cm$^2$). The pattern was then transferred to the PolySi using $SF_6/C_4F_8$ plasma etch as before.  The shallow regions were formed by a reactive ion etch (Oxford PlasmaPro80, 27mTorr, 175W, 35sccm Ar, 15 sccm $CHF_3$, DC bias ~448V, 25min). The final depth of the shallow region was measured to be 680 nm by a profilometer (KLA Tencor P15). The wafer was then diced (ADT proVectus 7100) and individual devices were bonded to fused silica coverslips (Esco Optics R425025) using low temperature sodium silicate bonding[3]. Before bonding, four through holes were drilled into the fabricated fused silica device using a 0.75mm triple ripple diamond drill bit (Crystalite) on a precision drill press (Servo Products Company 7170).

**Supplementary Note 1 Estimating the shot-noise limit by computer simulation**

To estimate the shot-noise limit, we simulated ideal FRET experiments where the underlying efficiency is constant and photon counting on the donor and acceptor channels follows Poisson statistics. We used realistic parameters extracted from experiments and the Gillespie algorithm for efficient simulation of the stochastic events of photon detection[4].

A simulated stream of donor and acceptor photons was generated in two steps. First, the inter photon arrival times were drawn from an exponential distribution parameterized by the total emission rate:

$$k_{tot} = k_{D,s} + k_{A,s} + k_{D,b} + k_{A,b}$$

where the first subscript denotes donor (D) or acceptor (A) channel and the second subscript specifies signal (s) or background (b). After this stream of photons was generated, the identity of each photon was then assigned (i.e. donor fluorescence, acceptor fluorescence, donor background, acceptor background) with probabilities given by the branching ratios. For example, the probability that a photon is signal from the donor channel is $k_{D,s} / k_{tot}$ .

A FRET histogram and measurement precision at N photons per efficiency calculation were then generated for this simulated photon stream following the procedures used for analyzing the experimental data.

**Supplementary Note 2 Estimating the shot-noise limit using the Cramér-Rao lower bound**

Here, we derive the Cramér-Rao lower bound (CRLB) for the variance of the maximum likelihood FRET efficiency estimator (MLE)[5]. From estimation theory, the CRLB sets the minimum variance of an unbiased estimator under given experimental conditions. In general, for a set of parameters to be estimated ($\hat{\theta}$),

$$\mathrm{cov}\left(\hat{\theta}\right) \geq CRLB\left(\theta\right) = F_{\theta}^{-1} \ , \tag{1}$$

where $F_{\boldsymbol{\theta}}$ is the Fisher information matrix derived from the likelihood function $\mathcal{L}\left(\theta \mid m\right)$. The elements of the Fisher information matrix are defined as

$$\{F_{\boldsymbol{\theta}}\}_{ij} = \left\langle \left(\frac{\partial}{\partial \theta_i} \ln L(\boldsymbol{\theta}|m)\right) \left(\frac{\partial}{\partial \theta_j} \ln L(\boldsymbol{\theta}|m)\right) \right\rangle_m \quad , \tag{2}$$

where the indices $i, j$ are over the elements in the set of parameters $\theta$, and the angled brackets indicate taking the expectation is over all possible values of the measurement, $m$.

Below we first derive the MLE for the FRET efficiency (E) from photon counting measurements and then we derive the analytical form of the CRLB.

In a photon-counting based smFRET experiment, the intensity (unit: # of photons per unit time) of the donor and acceptor channels can be expressed as

$$\begin{aligned} I_D &= I_T\left(1-E\right) + B_D \\ I_A &= I_T E + B_A \end{aligned} \ , \tag{3}$$

where $B_D$, $B_A$ are the background intensity on the donor and acceptor channels respectively and $I_T$ is the total intensity of the colorless stream from the FRET pair (background not included). We have omitted donor leakage ($\alpha$) in this treatment. With these definitions the probability of a given photon being on the donor ($p_D$) or acceptor ($p_A$) channel is,

$$
\begin{aligned}
p_D(E) &= \frac{I_D(E)}{I_D(E)+I_A(E)} \\
p_A(E) &= \frac{I_A(E)}{I_D(E)+I_A(E)} = 1 - p_D(E)
\end{aligned}
\quad . \tag{4}
$$

Consequently for N photons, the probability of detecting $n_D$ photons on the donor channel and $n_A$ photons on the acceptor channel is given by a Binomial distribution

$$
p\left(n_D, n_A \mid E\right) = \frac{N!}{n_A!n_D!}\left[p_A(E)\right]^{n_A}\left[p_D\left(E\right)\right]^{n_D} \quad . \tag{5}
$$

The likelihood function has the form

$$
\mathcal{L}\left(E \mid n_D, n_A\right) = p\left(n_D, n_A \mid E\right) \ , \tag{6}
$$

and the MLE of E is obtained from the condition

$$
\frac{d \ln \mathcal{L}}{dE} = 0 \ , \tag{7}
$$

which in our case gives

$$
\hat{E}_{MLE} = \frac{n_A I_T + n_A B_D - n_D B_A}{I_T\left(n_A + n_D\right)} \quad . \tag{8}
$$

Note that in the limit of zero background this reduces to the expected result:

$$
\hat{E}_{MLE} = \frac{n_A}{n_A + n_D} \quad . \tag{9}
$$

Moreover, by recognizing that

$$
\left\langle t_c \right\rangle = \frac{n_A + n_D}{I_T + B_A + B_D} \ , \tag{10}
$$

is the mean time to collect N total photons, it is straightforward to show that Eq.8 is equivalent to the common way of calculating FRET efficiency,

$$\hat{E}_{MLE} = \frac{n_A - B_A \langle t_c \rangle}{n_A - B_A \langle t_c \rangle + n_D - B_D \langle t_c \rangle} \ . \tag{11}$$

Given a likelihood function in the form of Eq. 5 and 6, the Fisher information can be calculated by applying Eq. 2. We then have

$$\begin{aligned} F(E) &= \left\langle \left( \frac{\partial}{\partial E} \ln \mathcal{L} \right)^2 \right\rangle_{n_A} \\ &= \frac{N I_T^2}{(E I_T + B_A)((1-E) I_T + B_D)} \end{aligned} \ , \tag{12}$$

where on the first line the expectation is taken over the possible values of the measurement (i.e. the possible values of $n_A$, since $n_D = N - n_A$). Note that to arrive at Eq. 12 we have taken advantage of the fact that $n_A$ follows Binomial statistics, which means that,

$$\langle n_A \rangle = N p_A(E) = N \frac{E I_T + B_A}{I_T + B_A + B_D}$$

$$\mathbf{var}(n_A) = N p_A(E)\big(1 - p_A(E)\big) = N \frac{(E I_T + B_A)\big((1-E) I_T + B_D\big)}{(I_T + B_A + B_D)^2} \tag{13}$$

Finally, the CRLB is

$$CRLB = F(E)^{-1} = \frac{(E I_T + B_A)\big((1-E) I_T + B_D\big)}{N I_T^2} \ . \tag{14}$$

Note that Eq. 14 reduces to the widely used formula $E(1-E)/N$ (Ref. [6,7]) in the absence of background. Figure S4 plots Eq.14 together with simulation, showing the excellent agreement between the two.

**Supplementary Note 3 Amplitude of the feedback forces on biomolecules**

The feedback electric field used in ABEL-FRET is in the range of $10^4$-$10^5$ V/m. Here we estimate the magnitude of the forces acting on the trapped molecule. A single element charge will feel a Coulomb force given by $F=eE$, which is in the 1-10fN range. On the other hand, a single molecule in aqueous solution experiences a "white" random force

$$\langle F(t_1)F(t_2)\rangle = g\delta(t_1-t_2)$$

The amplitude of this Langevin force is related to the viscous drag through the fluctuation-dissipation theorem[8]

$$g = 2\gamma k_B T$$

The amplitude of the force over a bandwidth $\Delta f$ is

$$\langle F\rangle_{\Delta f} = \sqrt{12\pi k_B T\eta a\Delta f}$$

where we have used the Stokes drag $\gamma=6\pi\eta a$. Using $a=5$ nm, $\Delta f=1000$ Hz (1ms time window) gives <F> ~20 fN. This is the magnitude of the random force that a single biomolecule experiences from random thermal collisions over a 1ms time window. Given that the feedback trap uses electrokinetic forces to compensate Brownian motion at ~1ms timescale, it is not surprising that the two forces are on the same order of magnitude. This simple analysis estimates the perturbation induced by electrokinetic forces to be very small (on the order of the Langevin forces).

A protein has both positive and negative residues that will feel forces along different directions. The energy scale of an electric dipole in a external field is defined by

$$E_{dp} = \left|\vec{p}\cdot\vec{E}\right|$$

where $\vec{p}$ is the intrinsic dipole moment of a protein. For this to reach $k_BT$, $p$ needs to be ~$1\times10^4$ debye. A recent computational survey[9] has shown that most protein chains (>99% surved) have dipole moments that are smaller than 2000 debye (mean = 543 debye). Thus the induced energy on a protein dipole is in general much smaller than the thermal energy. A similar argument was also given in Ref. [10] to show that the rotational dynamics of a single molecule are also not affected under the mild feedback electric fields in the ABEL trap.

**Supplementary Note 4 Assigning SSB binding stoichiometry based on diffusion coefficients**

It has been shown that the $D \sim (M)^{-1/3}$ scaling between the diffusion coefficient (D) of a protein and its molecular weight (M) approximately holds[11]. This relation predicts that ssDNA with two SSB tetramers bound would diffuse $\sim((M_{70}+M_{SSB}\times 2)/(M_{70}+M_{SSB}))^{-1/3}=0.83$ times slower than ssDNA with one SSB tetramer bound, where we used the known molecular weight of the $(dT)_{70}$ substrate ($M_{70}$ = 22kDa) and SSB tetramer ($M_{SSB}$ = 76kDa). In the experiments presented in Figure 4d and 4e of the maintext, the relative ratio of D between the low FRET state (E~0.45) and the high FRET state (E~0.93) is 0.82. We thus assign the low FRET state to be two SSB tetramers bound to $(dT)_{70}$ in the 35nt binding mode.

Further, to examine whether D of the high-FRET state is consistent with one SSB tetramer bound in the 65nt binding mode, we used the tool HydroPro[12] to estimate D of a bare SSB tetramer. HydroPro uses the atomic-level structure (PDB file) of the protein as input and considers the realistic shape of the protein in three dimensions to model hydrodynamic properties. Using a full-length SSB structure (pdb ID: 1eqq), HydroPro predicts a D value of 63.7μm$^2$/s. With a 22kDa ssDNA wrapped around in the $(SSB)_{65}$ mode, the complex is expected to have a $D= 63.7\times((22+76)/76)^{-1/3}=57$ μm$^2$/s. The measured D of the high-FRET state is 52 μm$^2$/s, consistent with our assignment.

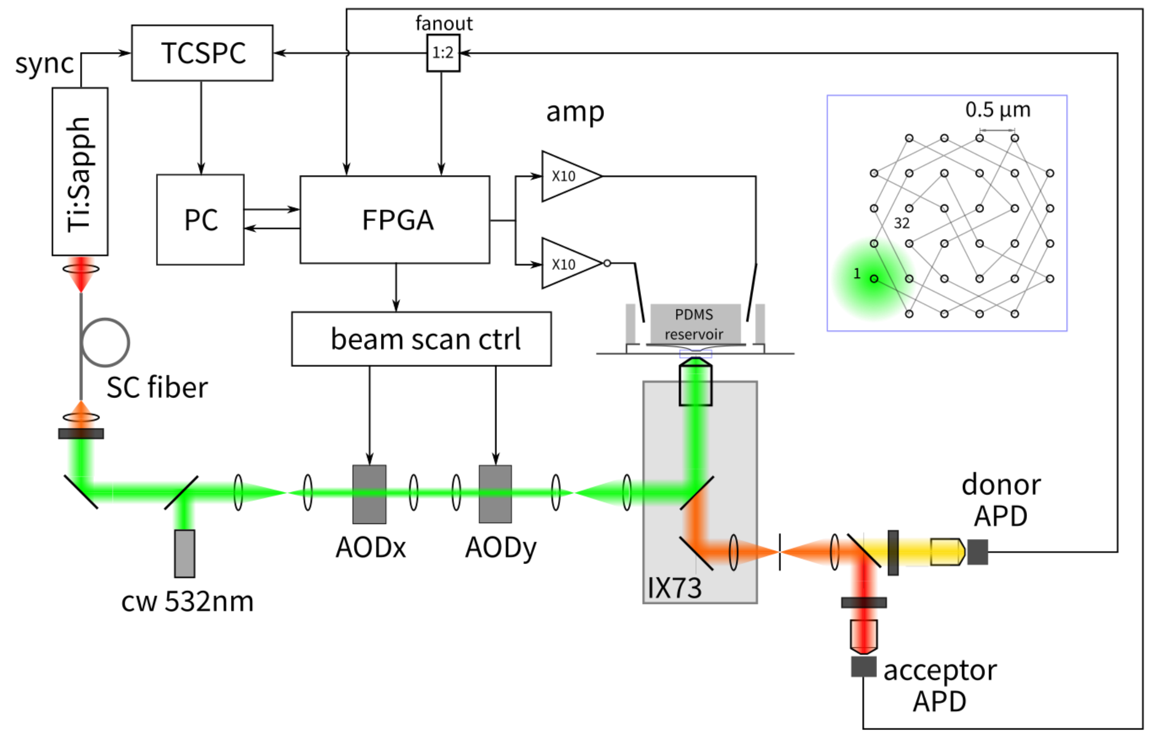


Figure S1 Detailed optical setup of ABEL-FRET. Ti:Sapph oscillator: Mira 900F (Coherent) pumped by a SproutD 10W (Lighthouse photonics). Supercontinuum (SC) fiber: gift from Robert Windeler, OFS Fitel, CW laser: Obis 532LS (Coherent). Acousto-optic deflector (AOD): MT110-B54A1,5-VIS (AA optoelectronics). AOD driver: DDSPA2X-D8b15b-34 (AA optoelectronics). Microscope base: Olympus IX73. Avalanche photodiode (APD): COUNT-T100 (Laser Components). TCSPC: Picoharp 300 (Picoquant). High-voltage amplifier: F10AD (FLC).

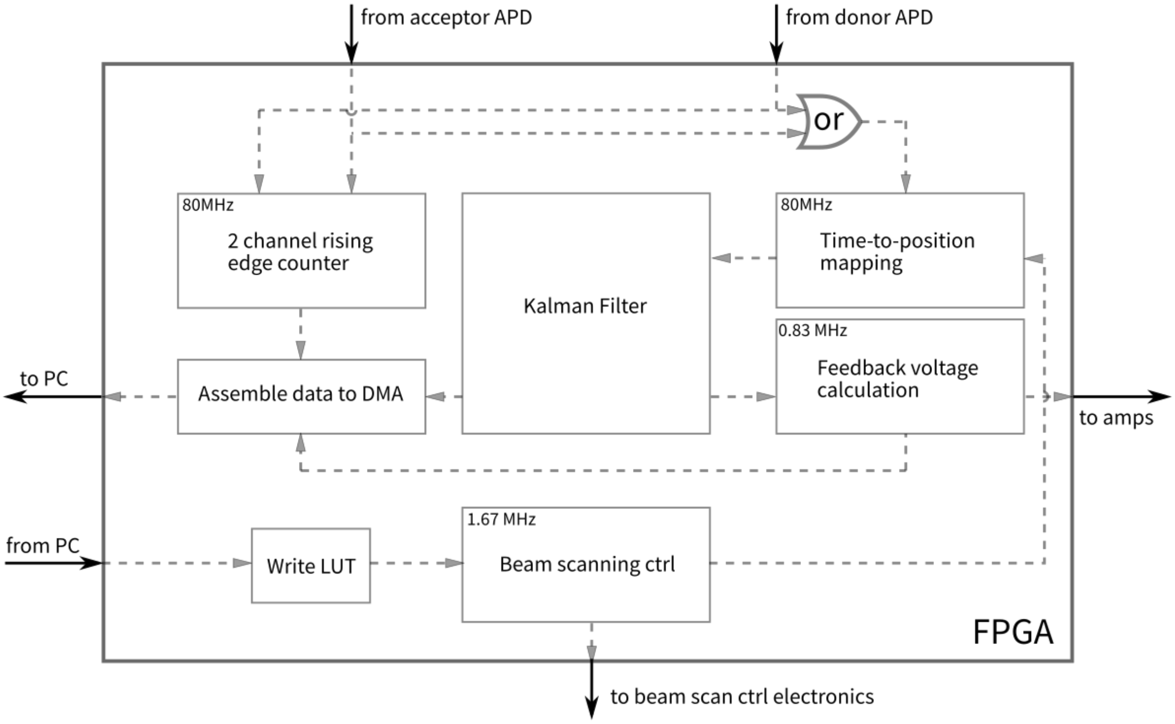


Figure S2 Logic schematic of the FPGA for real-time feedback control and dual-color photon counting.

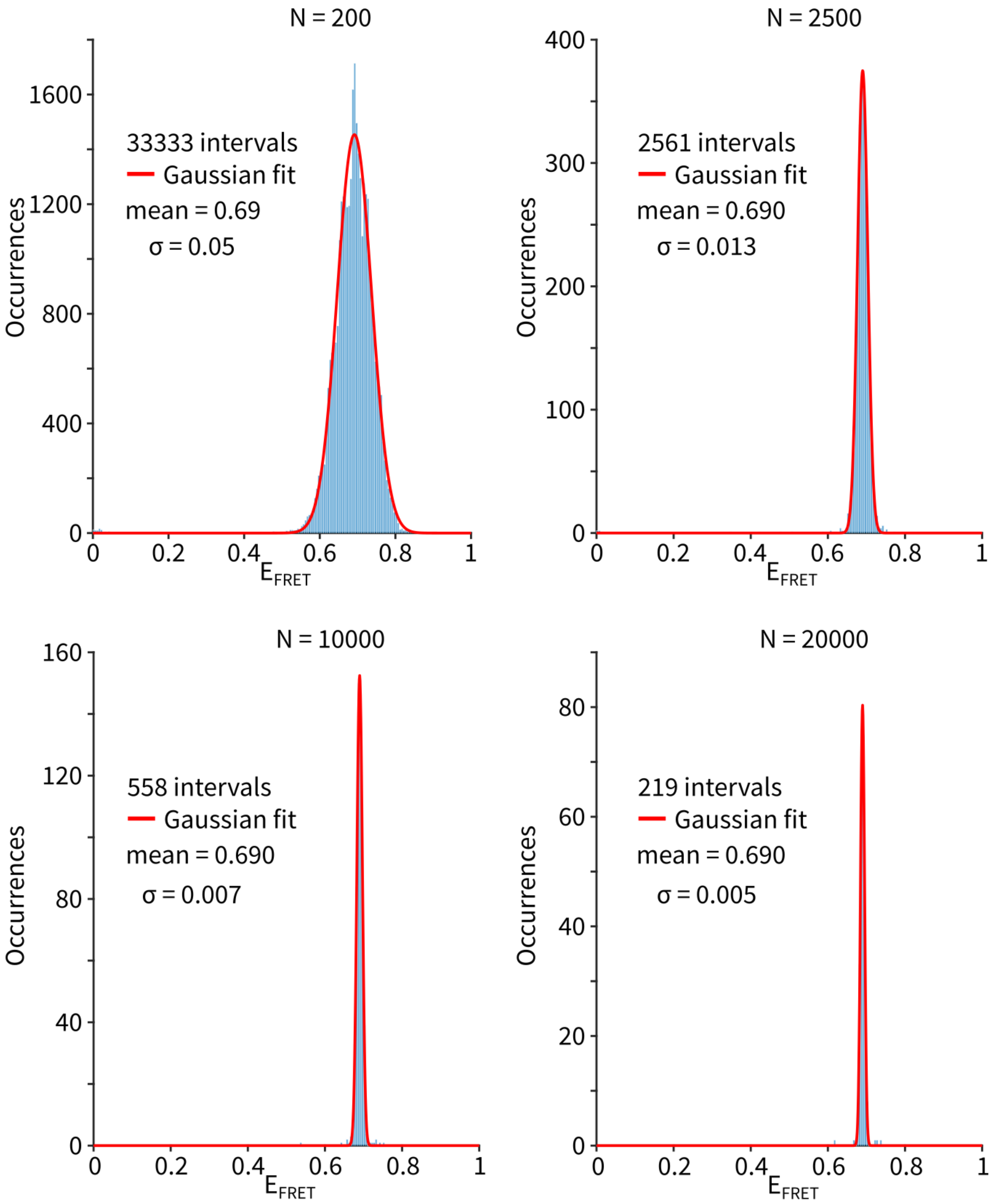


Figure S3 Individual FRET histograms and Gaussian fits of the 11bp dsDNA sample (a total of 226 molecules). The number of photons per efficiency calculation is marked at the top of the histogram.

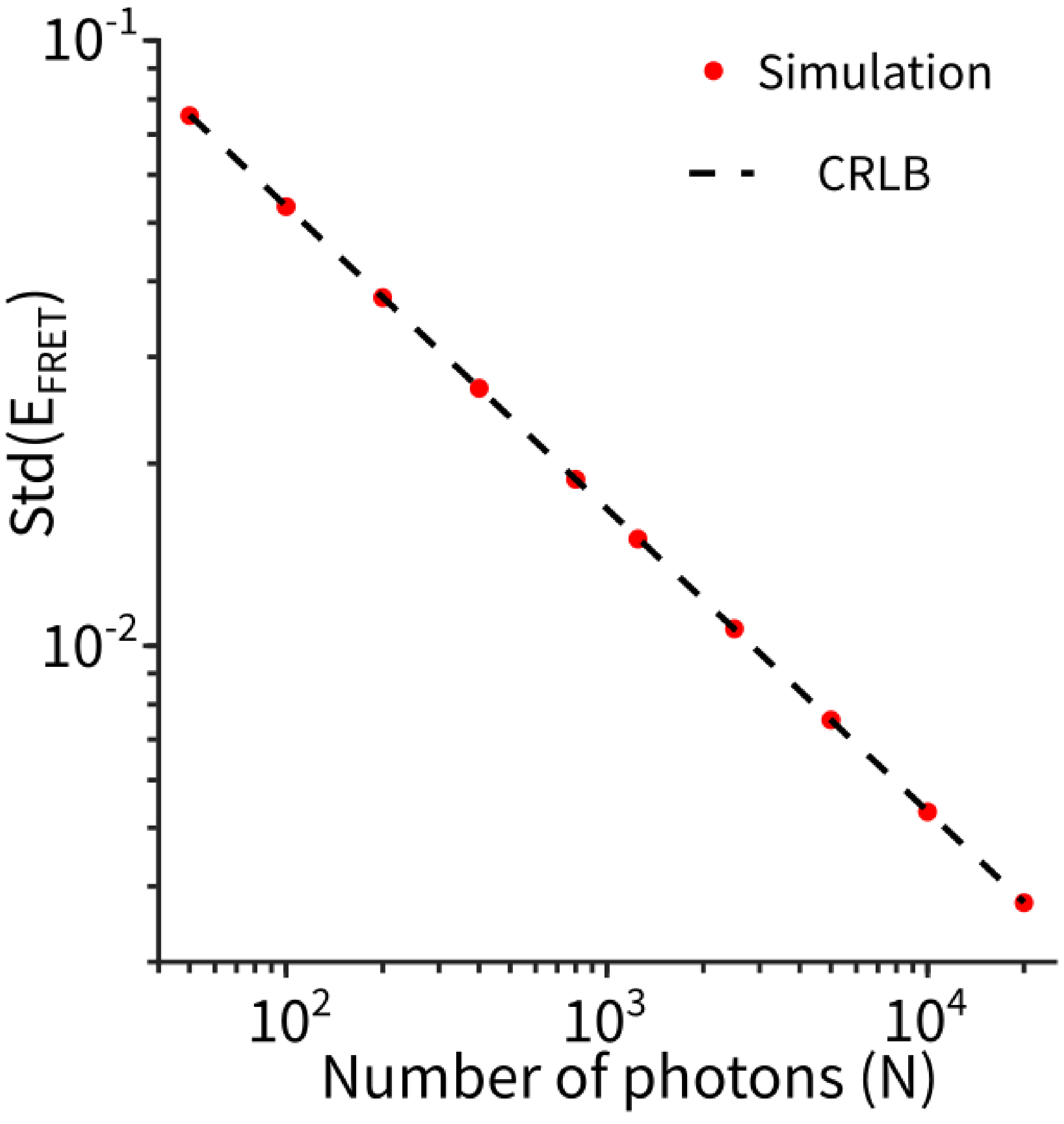


Figure S4 Simulation and Cramer-Rao bound calculation reach the same shot-noise limit of FRET efficiency.

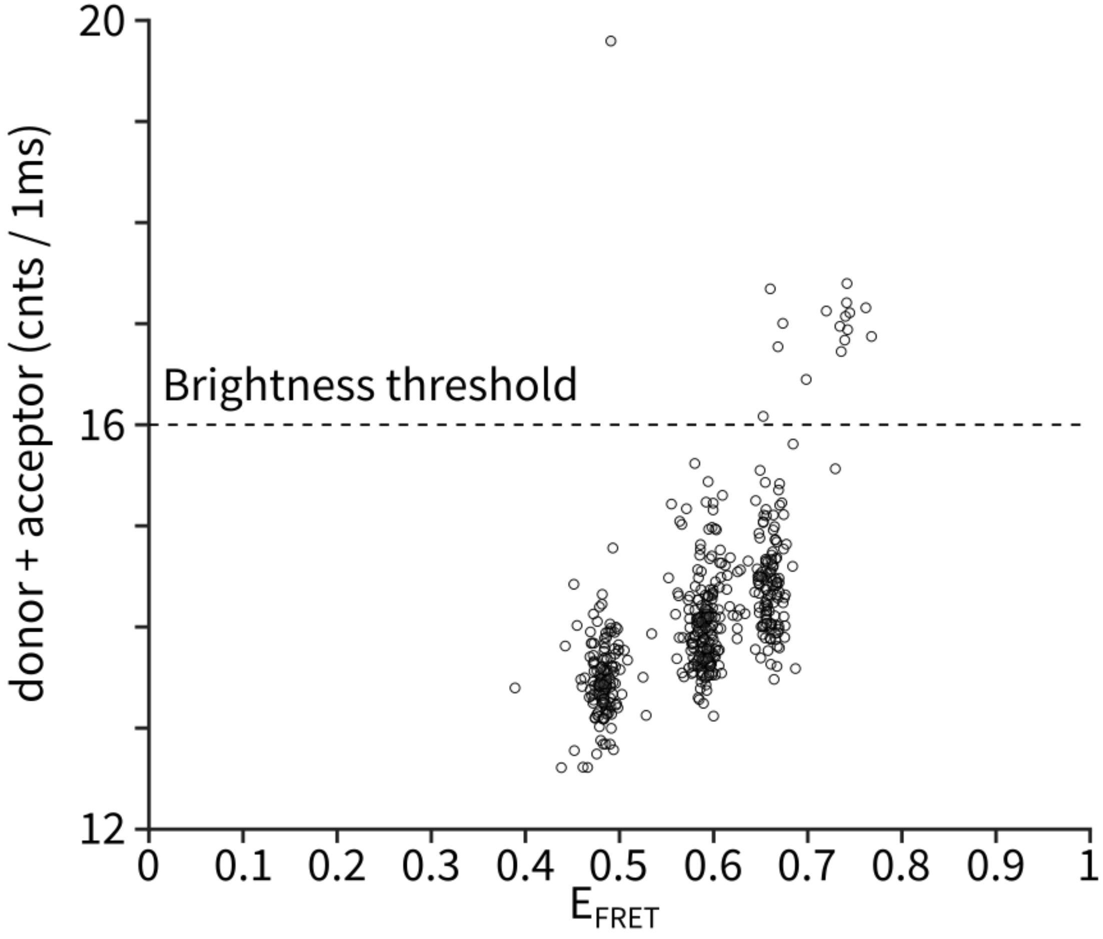


Figure S5 Resolving a mixture of 11, 12 and 13bp dsDNA molecules. 17 molecules (4%) are ~20% brighter than the rest of the population and are excluded in Figure 2c of the main text.

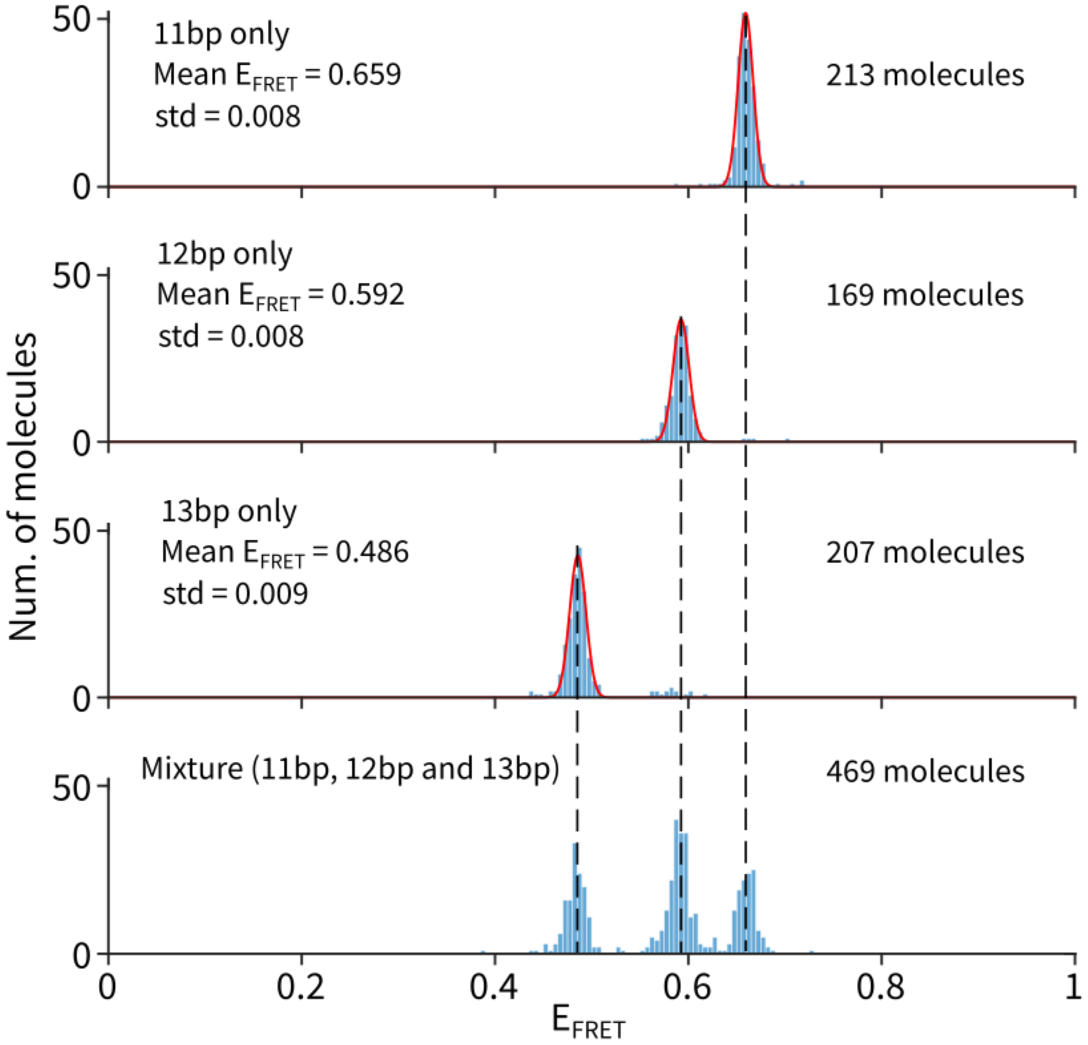


Figure S6 Identifiying the populations in the mixture sample by comparing to the samples of individual strands. Note that the efficiency of the 11bp sample (0.659) is slightly different than in Figure S3 (0.69). This is because the two experiments were done with different emission filters in the acceptor detection path.

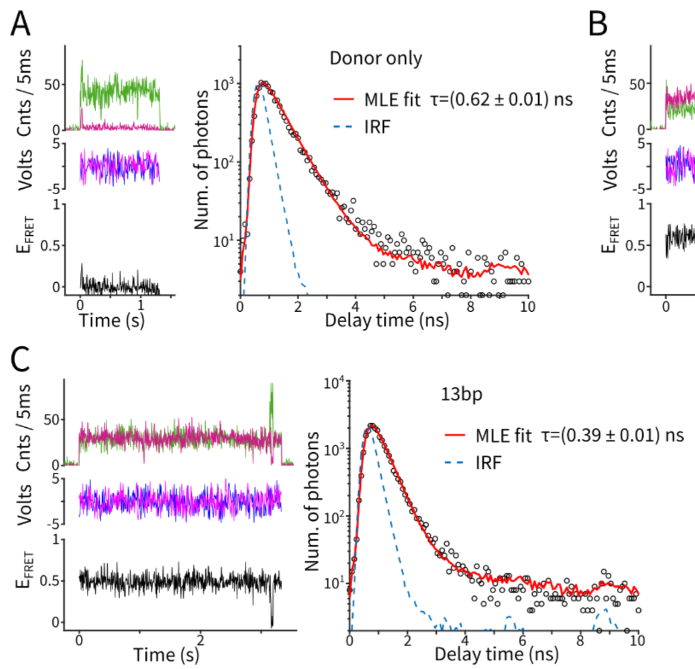


Figure S7 Simultaneous measurement of smFRET efficiency and donor excited-state lifetime. (**A-D**) Example single-molecule traces showing intensity, feedback voltages, FRET efficiency and photon delay time histograms.

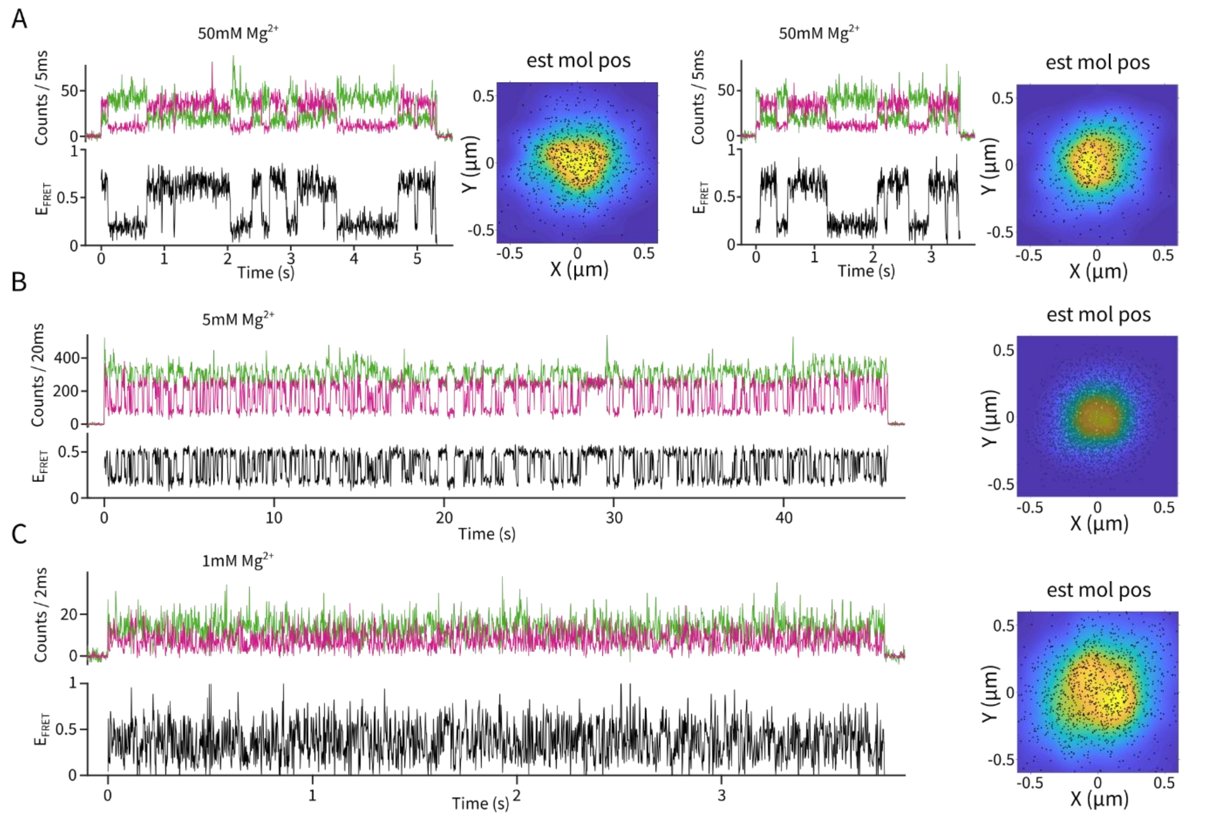


Figure S8 Example ABEL-FRET traces of Holliday junction dynamics. Here, instead of plotting the feedback voltages, we show the estimated molecule postion (down sampled every 100 points) during trapping to visualize the degree of residual motion of the molecule.

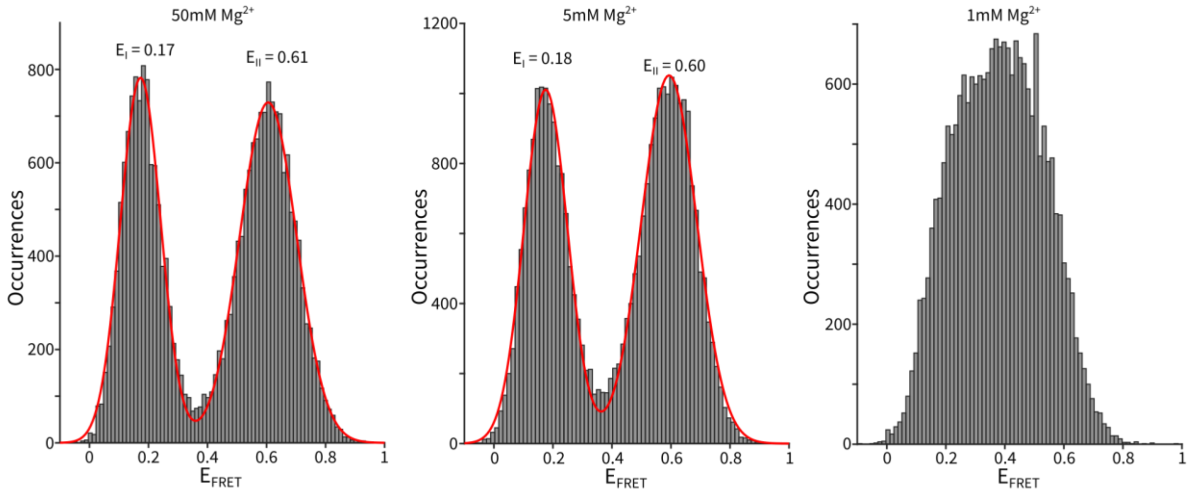


Figure S9 FRET efficiency histogram of Holliday junctions, calculated using the 5ms binned intensity data. In cases of 50mM and 5mM $Mg^{2+}$, the histogram was fit to a sum of two Gaussians. At 1mM $Mg^{2+}$, the dynamics are faster than the binning time. Number of molecules measured (from left to right): 62, 59 and 37.

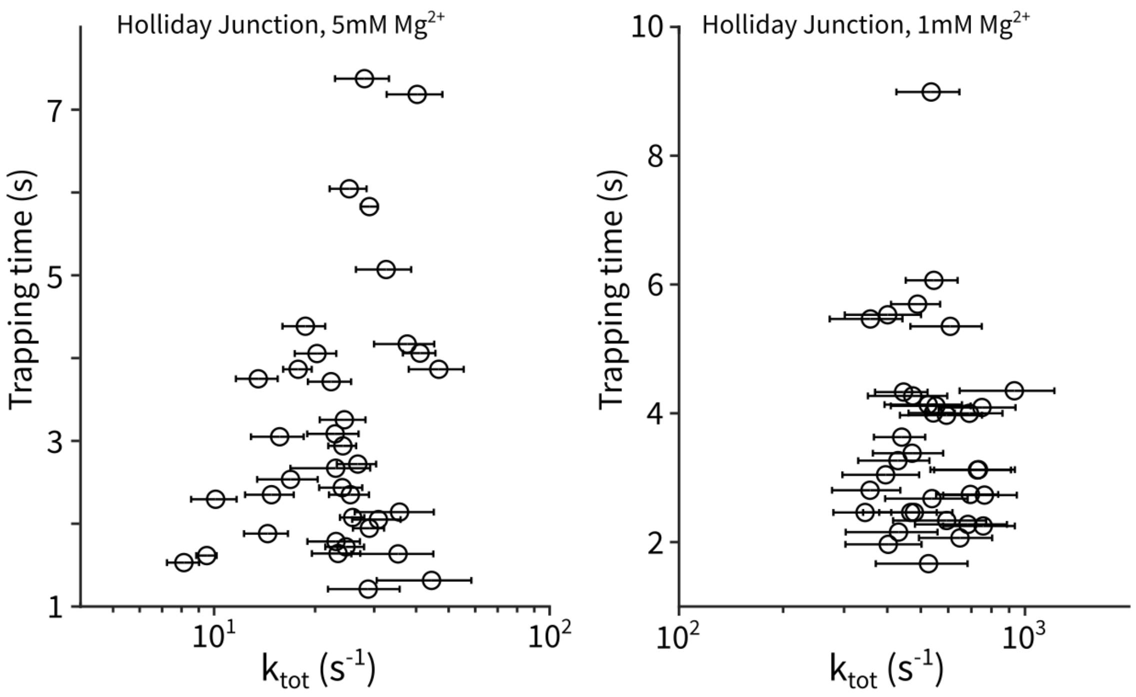


Figure S10 Revealing rate heterogeneity between single Holliday junction molecules. Every symbol represents a single molecule. Rates were determined by fitting the intensity cross-correlation with a single exponential. Error bars represent 95% confidence intervals from the fit.

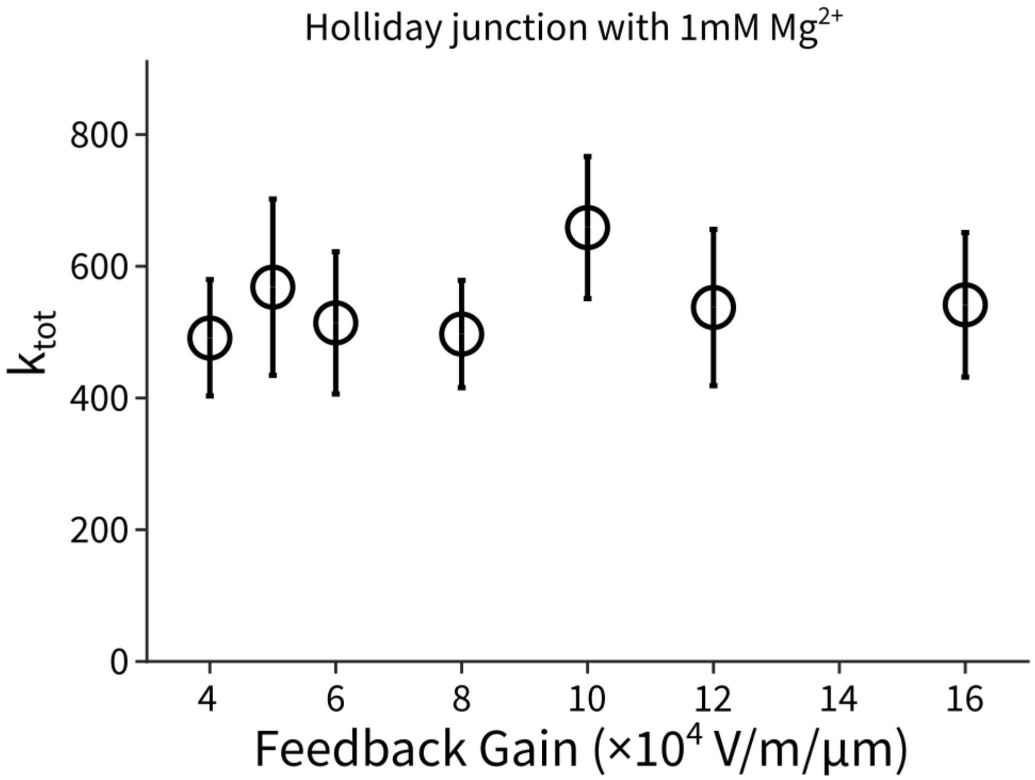


Figure S11 Holliday junction dynamics are not correlated with feedback voltage strength. The rates were determined by fitting the intensity cross-correlation averaged over many molecules. Error bars represent 95% confidence intervals from the fit. The number of molecules measured were (from low to high gain): 44, 31, 37, 31, 26, 13, 41.

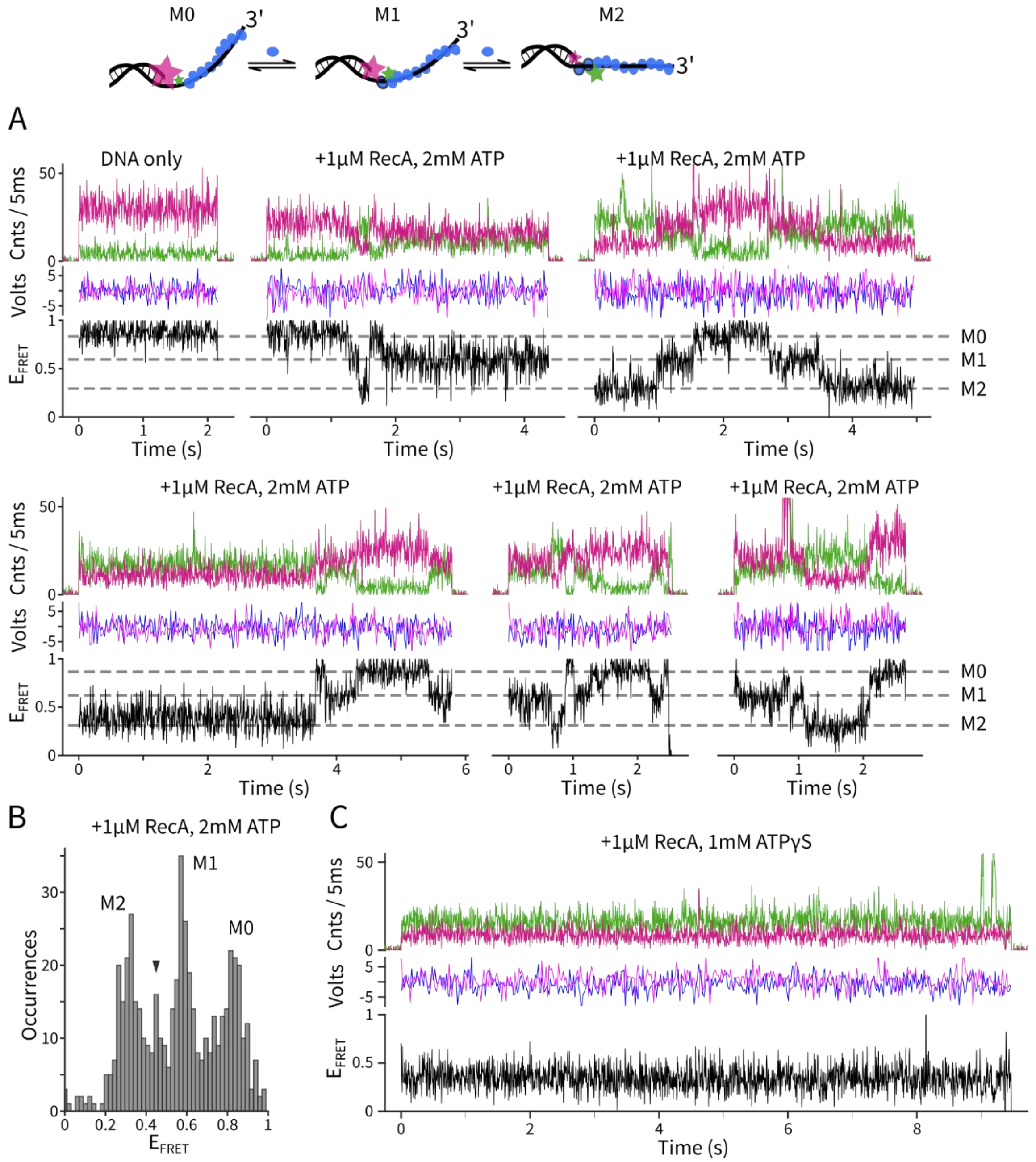


Figure S12 ABEL-FRET monitors RecA-ssDNA nucleofilament dynamics. (**A**) Example traces of trapped $(dT)_{10+49}$ molecules under the conditions marked above each panel. Dashed lines represent the three major FRET states. (**B**) Histogram of observed FRET states, showing three major populations at M0, M1 and M2 and at least one minor state (arrow) at ~0.45. (**C**) $(dT)_{10+49}$ in the presence of 1μM RecA and 1mM ATPγS only shows M2 state with no dynamics, representing fully formed nucleofilament.

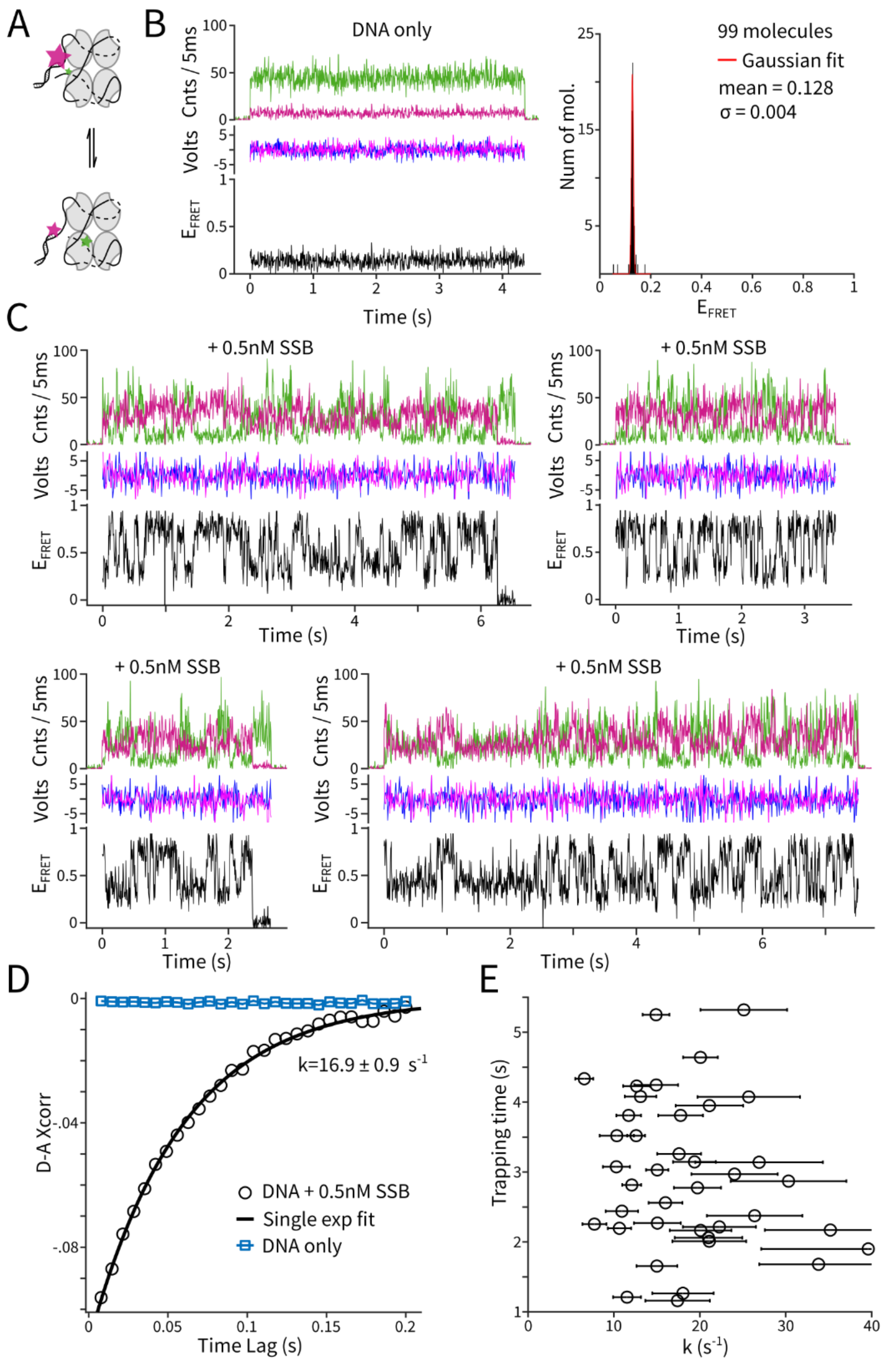


Figure S13 ABEL-FRET monitors SSB sliding on ssDNA in the 65 nucleotide binding mode. (**A**) Experimental design: the DNA substrate contains a 77nt ssDNA region and the FRET dyes are separated by 69nt. (**B**) Example traces and FRET efficiency histogram of DNA substrate only, without SSB. (**C**) Example traces in the presence of 0.5nM SSB with 200mM NaCl. (**D**) Intensity cross-correlation curves for DNA only (blue square, averaged over 99 molecules) and DNA with 0.5nM SSB (black circle, averaged over 40 molecules). Rate of SSB sliding was estimated by fitting with a single exponential function. (**E**) Revealing rate heterogeneity between single SSB proteins. Every symbol represents a single molecule. Error bars represent 95% confidence intervals from the fit.

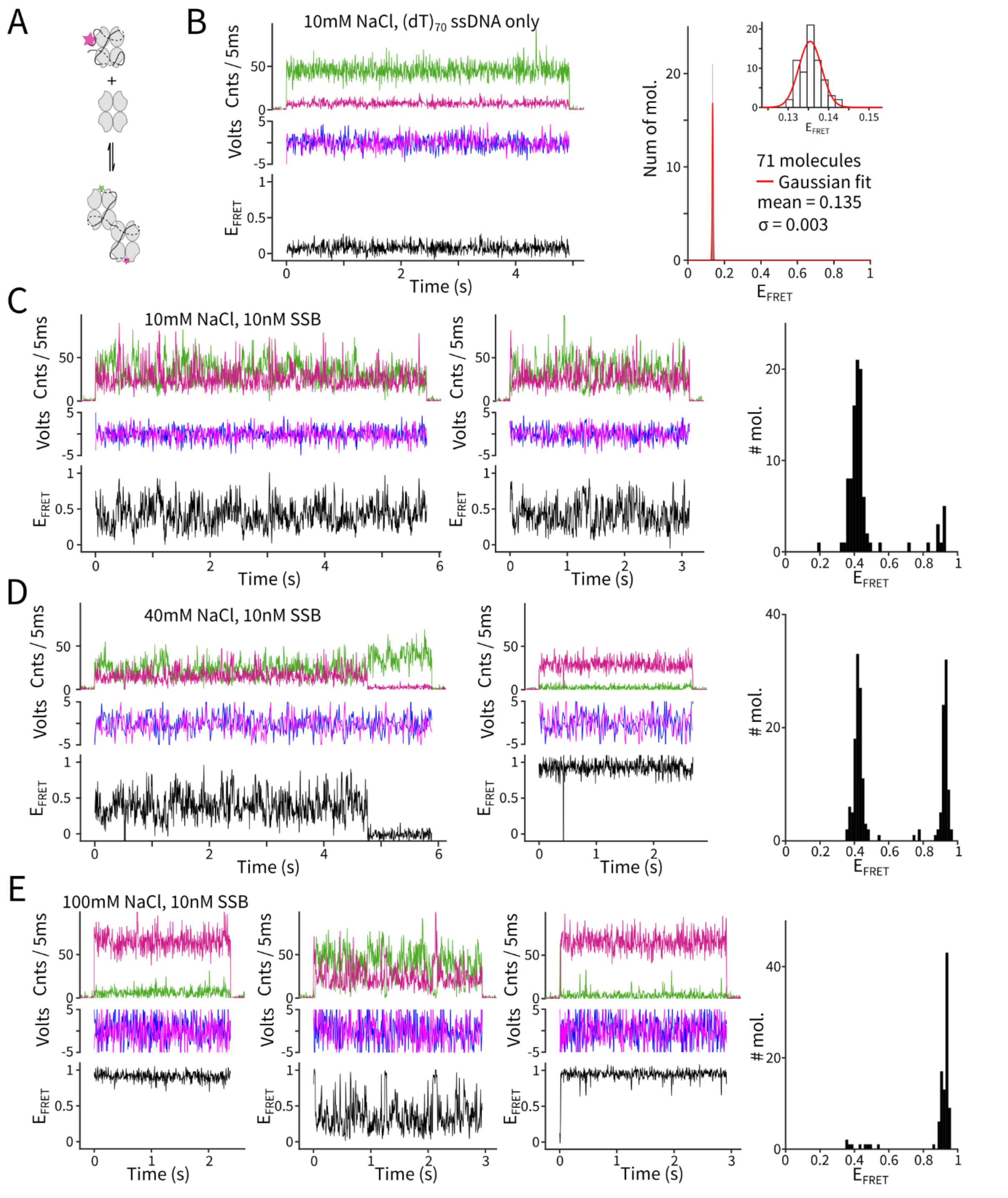


Figure S14 ABEL-FRET monitors SSB binding modes. (**A**) Experimental design: the DNA substrate is Cy3 and Cy5 end-labeled $(dT)_{70}$ ssDNA. In the 65nt binding mode, the ends of the ssDNA are brought to close proximity, resulting in high FRET. In the 35nt binding mode, the $(dT)_{70}$ can bind two SSB tetramers and the ends of the ssDNA are further apart, giving a lower FRET signal. (**B**) Example traces and FRET efficiency histogram of DNA substrate only, without SSB. (**C-E**) Left: example traces with DNA substrate and SSB at different NaCl concentrations. Right: histograms of FRET efficiency.

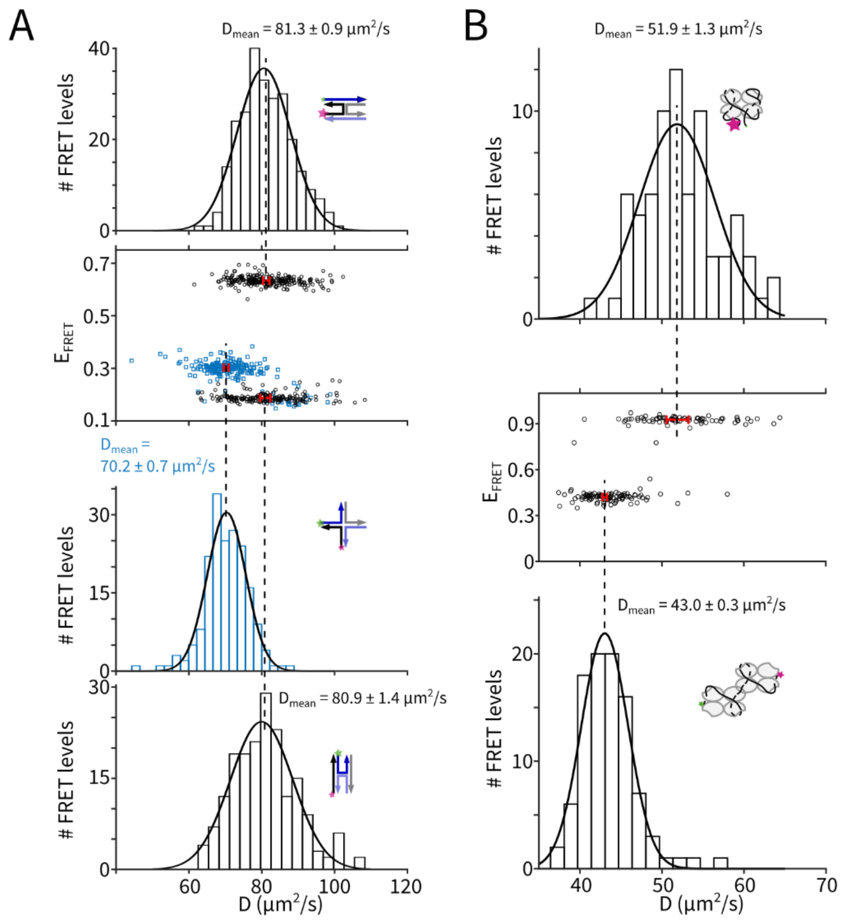


Figure S15 Hydrodynamic profiling of HJ and SSB-ssDNA complexes. (**A**) Diffusion coefficient histograms of the FRET-separated populations of HJ. Solid curves are Gaussian fits. (**B**) Diffusion coefficient histograms of the FRET-separated populations of SSB-ssDNA complexes. Numbers indicate mean values of the D histograms with 95% confidence intervals.

Table S1 Parameters used in simulation (Supplementary Note 1) and Cramér-Rao bound calculations (Supplementary Note 2).

| Parameter | Value |
| --- | --- |
| E | 0.69 |
| $k_{D,s}+k_{A,s}$ | 15000 $s^{-1}$ |
| $k_{D,b}$ | 1400 $s^{-1}$ |
| $k_{A,b}$ | 230 $s^{-1}$ |

Table S2: Comparison of ABEL-FRET with other smFRET modalities

| Technique | Analyte limitations | Maximum Observation time | Time Scale | Throughput | Excited-state lifetime | Measure D | Ref |
|---|---|---|---|---|---|---|---|
| ABEL-FRET | - | ~10s | ns-10s | ~600 mol/h | Yes | Yes (SM*) | This work |
| Burst based confocal | - | ~1ms | ns-ms | ~3000-12000 mol/h[13] | Yes | Yes (ensemble) | [14–17] |
| TIRF-FRET | Requires tethering | >100s | 10ms-100s | 1000 per FOV (EMCCD) ~15000 per FOV (sCMOS)[18] | No | No | [19–21] |
| Surface confocal | Requires tethering | >100s | ns-100s | <1000 overnight[22] | Yes | No | [22,23] |
| SWIFT | D < 45 $\mu m^2/s$ | ~10s | 10ms – 20s | ~2000/ min Only 2% observed longer than 5s for 201bp dsDNA | No | Possible (SM) | [24] |
| Piezo-tracking FRET | D < 0.5 $\mu m^2/s$ | <1s (for dsDNA in 90% glycerol) | ns-s | ~100s mol/h† | Possible | Possible | [25] |
| Capillary cell | - | ~5ms-1s (flow rate dependent) | 0.1ms-s | ~1000 mol/h† | No | Possible | [26,27] |
| Dimple Machine | - | > 100s | 10ms-100s | 2000 mol/h[28] | No | No | [29] |
| Liposome tethering | Tether to liposome | 10-50ms | ns-10ms | ~1000 mol/h† | Possible | No | [30] |
| Single-molecule recycling | - | ms per snapshot ~10s total | <ms | NA | Yes | Yes (ensemble) | [31] |

† Estimated based on operation principle   * SM = single-molecule

Table S3: DNA sequences

| Experiment | Sequences* |
|---|---|
| Short dsDNA strands† | 11bp:<br>5'-GAG TAC ACG TG-**Cy3** -3'<br>5'-CAC GTG TAC TC-s*Cy5*-3'<br><br>12bp:<br>5'- GAG TAG CAC GTG-**Cy3** -3'<br>5'- CAC GTG CTA CTC–s*Cy5* -3'<br><br>13bp:<br>5'- GAG TAG ACA CGT G-**Cy3** -3'<br>5'- CAC GTG TCT ACT C-s*Cy5*-3' |
| Holliday junction (Ref.[32]) | J7b: 5'-**Cy5**-CCC TAG CAA GCC GCT GCT ACG G-3’<br>J7h: 5'-**Cy3**-CCG TAG CAG CGC GAG CGG TGG G-3’<br>J7r: 5'-CCC ACC GCT CGG CTC AAC TGG G-3’<br>J7x: 5'-CCC AGT TGA GCG CTT GCT AGG G-3’ |
| $(SSB)_{65}$ diffusion $(dT)_{69+8}$ (Ref.[33]) | 5'-TGG CGA CGG CAG CGA GGC TTT TTT TTT TTT TTT TTT TTT TTT TTT TTT TTT TTT TTT TTT TTT TTT TTT TTT TTT TTT TTT TTT TTTsCy3TTT TTT TT-3'<br><br>5’-**Cy5**-GCC TCG CTG CCG TCG CCA |
| SSB binding mode $(dT)_{70}$ | 5’-**Cy3** TTT TTT TTT TTT TTT TTT TTT TTT TTT TTT TTT TTT TTT TTT TTT TTT TTT TTT TTT TTT TTT TTT TTT TTT T-s*Cy5*-3’ |
| RecA binding $(dT)_{10+49}$ (Ref. [34]) | 5'-TGG CGA CGG CAG CGA GGC TTT TTT TTT TTCy3T TTT TTT TTT TTT TTT TTT TTT TTT TTT TTT TTT TTT TTT TTT TTT TTT-3'<br><br>5'-**Cy5**-GCC TCG CTG CCG TCG CCA -3' |

* **Bold**: incorporated during synthesis (/3Cy3Sp/, /3Cy5Sp/, /5Cy5/ or /5Cy3/)

Underscore: Amino Modifier C6 dT, reacted with Sulfonated Cy3 or Cy5-NHS (sCy3 or sCy5)

*Italic*: 3’-amino modification(/3AmMO/), reacted with Sulfonated Cy3 or Cy5-NHS (sCy3 or sCy5)

†Extra bases were added in the middle of the strands to preserve the environment around the dyes at the ends

Table S4: Sources of reagents

| Reagent | Source |
|---|---|
| Cyanine3-NHS | Lumiprobe 11320 |
| Cyanine5-NHS | Lumiprobe 13320 |
| mPEG-silane | Gelest SIM6492.73 or Laysan Bio MPEG-SIL-5000-1g |
| HEPES | Sigma, H4034 |
| 5M NaCl | Ambion AM9760G |
| 1M $MgCl_2$ | Ambion AM9530G |
| ATP | Roche 10519979001 |
| ATPγS | Roche 11162306001 |
| Trolox | Sigma-Aldrich 238813 |
| Protocatechuate-3,4-dioxygenase | Oriental Yeast Co., 46852004 |
| Protocatechuic acid | Sigma-Aldrich 37580 |
| E coli. RecA | New England Biolabs M0249L |
| E coli. SSB | Promega M3011 |
